# Molecular Mechanocytometry Using Tension-activated Cell Tagging (TaCT)

**DOI:** 10.1101/2023.01.10.523449

**Authors:** Rong Ma, Arventh Velusamy, Sk Aysha Rashid, Brendan R. Deal, Wenchun Chen, Brian Petrich, Renhao Li, Khalid Salaita

**Author notes:** These authors contributed equally to this work (Arventh Velusamy, Sk Aysha Rashid, Brendan R. Deal).

## Abstract

Flow cytometry is routinely used to measure single-cell gene expression by staining cells with fluorescent antibodies and nucleic acids. Here we present Tension-activated Cell Tagging (TaCT) to fluorescently label cells based on the magnitude of molecular force transmitted through cell adhesion receptors. As a proof-of-concept, we analyzed fibroblasts and mouse platelets after TaCT using conventional flow cytometry.

---

Using antibodies and complementary nucleic acids to fluorescently label cells followed by analysis and sorting using flow cytometry and Fluorescence-activated Cell Sorting (FACS) is a cornerstone of modern biological and biomedical sciences. Flow cytometry is typically used for characterizing single-cell molecular expression levels, i.e., the biochemical profile of cells. The mechanical profile of cells offers a complementary *biophysical* marker that is intimately coupled to their biochemical profile.[1-3] Accordingly, a suite of flow cytometry methods have been developed to readout cell mechanics by measuring cell deformation during hydrodynamic stretching or micro-constriction.[4-6] For example, in real-time deformability cytometry, a high-speed camera records the transient deformation of cells while passing through a narrow channel under shear flow.[7] Deformation-based cytometry methods are powerful and have demonstrated clinical potential by detecting disease-specific changes in the mechanical deformation of cells.[8, 9] However, one major challenge is that cell deformation is a very convoluted product of multiple parameters that include cell volume, membrane composition, cytoskeletal dynamics, and cell cycle.[10-13] Thus, it would be highly desirable to develop mechanophenotyping platforms that employ molecular markers of mechanotransduction. To date, the readout of molecular forces generated by cells is performed using high-resolution fluorescence microscopy, which is low throughput. Indeed, there are no flow cytometry or FACS-like techniques that can tag cells based on the magnitude of molecular forces generated (**Table S1**).[14, 15]

To address this gap, we present Tension-activated Cell Tagging (TaCT) (**Figure 1A)**, which enables flow cytometry-based identification and sorting of mechanically active cells based on the molecular forces transmitted by their surface adhesion receptors. TaCT probes are engineered DNA duplexes that have a digital response to pN force and release a cholesterol-modified strand that spontaneously incorporates into the membrane of force-generating cells.

**Figure 1.**
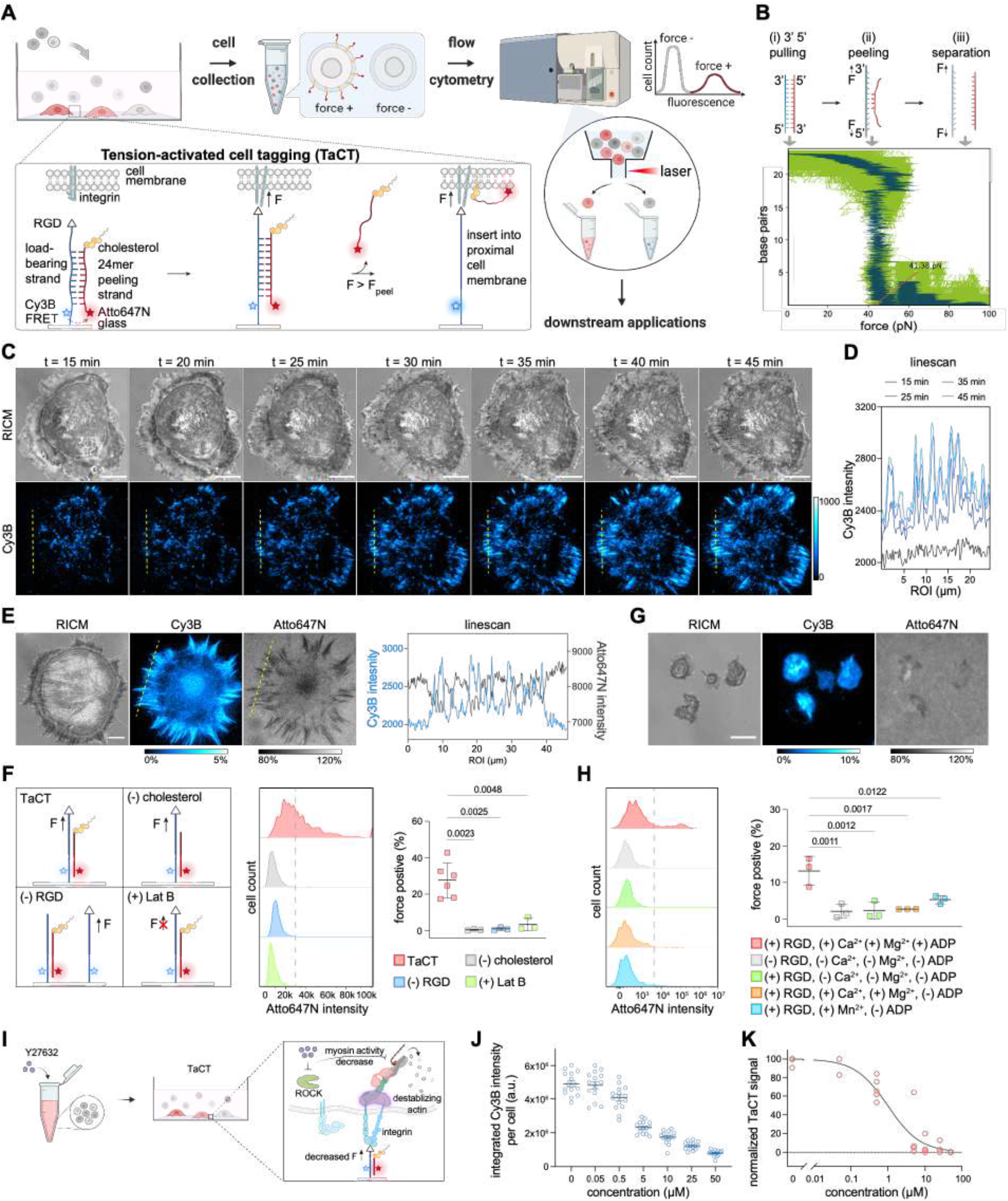
Tension-activated cell tagging (TaCT). (A) Schematic showing TaCT assay. (B) TaCT is based on 3’5’ mechanical pulling of a DNA duplex that leads to its dehybridization. Plot of 24mer dsDNA stability as a function of applied force generated using oxDNA simulation with a loading rate of 2.81×10^3^ nm/s (shown light green). Dark green shows the exponential moving average (EMA) and indicates a 41 pN dehybridization transition. (C) Time-lapse reflection interference contrast microscopy (RICM) and Cy3B fluorescence images of NIH3T3 cell spreading on TaCT surface 15-45 min after seeding. Scale bar = 10 µm. (D) Linescan of Cy3B images noted by yellow dashed line in (C). (E) Representative RICM, Cy3B, and Atto647N microscopy images of NIH3T3 cell cultured on TaCT substrate. Intensity bar for Cy3B image indicates the % of TaCT probes that undergo force-triggered release while the intensity bar for Atto647N shows the signal normalized to the background of intact TaCT probes. Scale bar = 10 µm. Linescan shows anti-colocalization of Cy3B and Atto647N intensities. (F) Flow cytometry histograms of NIH3T3 cells cultured on TaCT substrates as well as control surfaces with (-)cholesterol, (-)RGD, and (+)Lat B. Plot displays the force positive population from n=3-6 biological replicates (mean±SD, two-tailed student’s t-test). Grey dashed line indicates the gate for determining force-positive population. (G) Representative RICM, Cy3B, and Atto647N microscopy images of mouse platelets on TaCT substrates. Scale bar = 5 µm. (H) Representative flow cytometry histograms of mouse platelets incubated on TaCT substrates. Grey dashed line indicates the gate for determining force-positive population. Controls included experiments with TaCT probes -RGD (grey), -divalent cations -ADP agonist (green), -ADP agonist (orange), and -ADP/+Mn^2+^ (blue). Data plotted from n=3 animals (mean±SD, two-tailed student’s t-test). (I) Schematic showing Y27632 treatment decreases integrin forces. (J) Quantitative analysis of the integrated Cy3B signal per cell from microscopy images of MEF cells pretreated with 0, 0.05, 0.5, 5, 10, 25, and 50 µM of Y27632 and cultured on peeling probe substrate. Plot shows the result from n=3 biological replicates (mean±SEM). (K) TaCT signal of cells pretreated with different concentration of Y27632. Plot shows the result from n=2-4 biological replicates.

TaCT takes advantage of the fundamental mechanism of double stranded DNA “peeling” under force.[16] When a short DNA duplex is stretched from both ends of one of the strands (**Figure 1B**, 3’-5’ pulling), the duplex is destabilized, and rapidly denatures with sufficient force **(Figure 1B**, peeling), leading to the complementary DNA separation (**Figure 1B**, iii. separation).[16-18] Due to the narrow range of forces at which the denaturation transition occurs (**Figure 1B**, ii. peeling, **Figure S1**), force-induced peeling can be characterized as a simple two-state system, i.e., we can treat the DNA duplex as either being in dsDNA form or in completely separated ssDNA form. For example, a 24mer duplex (sequence in **Table S2**) was reported to peel at *F* = 41 pN using magnetic tweezer measurements.[16] To confirm the critical peeling force for this 24mer, we used the oxDNA coarse-grained model to apply a load of 2.81×10^3^ nm/s and recorded the number of base pairs in the duplex.[19] The peeling force (*F*_peel_) was defined as the force at which the number of base pairs in the duplex is ≤5. We found that *F*_peel_ = 41±2.8 pN (mean±SD) by averaging the first 100 data points in the simulation (**Figure 1B, Video S1, Figure S1**).

The TaCT probe was comprised of a load-bearing strand and a 24mer peeling strand. The load-bearing strand displayed the RGD integrin ligand at one terminus and was attached to the glass slide through its other terminus. The loading-bearing strand also incorporated an internal Cy3B dye. The complementary peeling strand was designed to release once *F*>*F*_peel_ and its termini were labeled with Atto647N and cholesterol (**Figure 1A, Figure S2**). The TaCT probes were annealed and immobilized on streptavidin-coated glass slides (**Figure S3**). At rest, the probe is in the dsDNA form where Cy3B and Atto647N form a FRET pair. When *F*>*F*_peel_, the Atto647N strand peels and dissociates, while the Cy3B dye on the load bearing strand is de-quenched. The FRET efficiency for TaCT probes was 93.8% and hence the Cy3B signal is enhanced by >10 fold upon peeling (**Figure S4A**). The probe density was measured at 5200±286 molecules/µm^2^ using a quantitative fluorescence calibration, which then allowed us to convert the Cy3B fluorescence signal to %peel (**Figure S4B, C**).[20, 21]

To demonstrate that cell-generated forces drive DNA duplex peeling, mouse embryonic fibroblasts (MEF) were plated on TaCT probe surfaces and then imaged using conventional fluorescence microscopy (**Figure 1C, Video S2**). A timelapse video showed cell spreading that coincided with Cy3B signal, confirming integrin binding and engagement with RGD ligands on the load-bearing strands. The Cy3B signal was primarily localized to the cell edge, consistent with the distribution of focal adhesions (FAs), and accumulated as a function of time (**Figure 1D**). The growth of Cy3B signal corresponded to the loss of Atto647N signal and confirmed the force-induced peeling mechanism (**Figure 1E**). This mechanism can generate high quality maps of integrins forces independent of cholesterol conjugation to the probe (**Extended Figure 1A, Figure S5**). Quantitative microscopy analysis of 227 cells showed that 0.9±0.3% of probes peeled under each cell, equivalent to ∼47±16 mechanical events/µm^2^ with *F*>41 pN (**Extended Figure 1B**). The observed tension signal colocalized with markers of FAs such as vinculin and phosphorylated focal adhesion kinase (FAK pY397), as well as actin stress fibers (**Extended Figure 1C, D, E**), confirming that forces were primarily transmitted by integrins within FAs.[22] Importantly, the peeling signal mirrors that of the turn-on tension gauge tether (TGT) probes (**Extended Figure 2A, B**) but avoids the termination of mechanotransduction, which represents a major advantage as a molecular force sensor (**Extended Figure 2A, C, D, Supplementary Note**). Control groups of cells treated with Latrunculin B (Lat B), which inhibits actin polymerization and disrupts force generation, showed significantly reduced peeling signal, as expected (**Extended Figure 3A, B)**. Cells treated with Lat B 50 min after seeding did not show tension signal changes, confirming that the peeling mechanism is irreversible, and maps accumulated mechanical events over time. As expected, addition of soluble peeling strand led to the loss of the tension signal in cells treated with Lat B, showing a simple approach to resetting tension signal and recording real-time events (**Extended Figure 3C, D, E, F**).

After confirming that DNA peeling faithfully maps molecular traction forces, we next investigated cholesterol-DNA cell tagging for TaCT (**Figure 1A**). Prior work showed that cholesterol-DNA can partition into the plasma membrane of cells.[23, 24] We discovered that cholesterol-ssDNA conjugates are ∼50 fold more effective at tagging cells compared to cholesterol-dsDNA conjugates (**Extended Figure 4A, B, Figure S6**). This is advantageous as it enhances the specificity of TaCT. We also verified that cholesterol-ssDNA membrane association is linearly proportional to the soluble conjugate concentrations tested (**Extended Figure 4C, D**).[24] The stability of TaCT tags was examined by incubating cholesterol-ssDNA with cells, washing, and measuring the loss of DNA as a function of time. We found that ∼20-30% of the cholesterol-tethered DNA dissociated at 90 min (**Extended Figure 4E, F**). Accordingly, we chose to incubate the cells on the surface for 1 h, which allowed for focal adhesion formation and TaCT to proceed while limiting cholesterol dissociation.

As a proof-of-concept of TaCT, NIH3T3 cells were seeded onto the TaCT substrates, allowed to spread and then collected by gentle scraping. TaCT signal was then measured by flow cytometry. Probes lacking RGD, probes lacking cholesterol as well as full TaCT probes but with cells treated with Lat B were used as negative controls. The resulting flow cytometry histograms showed a heterogeneous distribution of TaCT signal, matching the tension signal distributions observed under microscopy (**Figure 1F, Extended Figure 1B**). The cells on the TaCT substrate showed 27.6 ±4.0% force positive population compared to 0.6 ±0.2%, 1.1 ± 0.5%, and 3.5 ± 2.0% in (-)cholesterol, (-)RGD, and (+)Lat B controls, respectively (**Figure 1F, Extended Figure 5**). The Atto647N geometric mean fluorescence intensity (gMFI) of cells incubated on TaCT showed a >2-fold increase compared to controls, whereas the Cy3B gMFI did not change (**Extended Figure 5**). Taken together, this result confirmed that TaCT is triggered by force transmission through the integrin-RGD complex, followed by cholesterol-mediated membrane incorporation. Background tagging of cells due to cell spreading and proximity to the surface leads to negligible TaCT signal (**Extended Figure 6**).

Next, we tested whether this strategy works in primary cells. Platelets were chosen for this demonstration as mechanical forces play a crucial role in platelet activation, aggregation, and clot retraction, which are necessary steps in coagulation.[25] When mouse platelets were seeded onto the TaCT surface for 30 min, we noted loss of Atto647N signal that was coupled with an increase in Cy3B. This confirms that platelet integrin force transmission was sufficient to mediate DNA peeling with *F*>41 pN (**Figure 1G**). The Atto647N depletion signal did not colocalize exclusively with the Cy3B turn-on signal, which may be due to the accumulation of cholesterol-DNA probes at the basal face of the cell membrane **(Extended Figure 7A, B)**. Nonetheless, the TaCT signal was measured by flow cytometry, which showed 13.3±2% force positive population (**Figure 1H, Extended figure 7C**). Control experiments withholding the divalent cations Mg^2+^ or Ca^2+^ necessary for full integrin activation showed minimal TaCT signal and force positive population. Likewise, withholding ADP agonist inhibited platelet forces generation and TaCT signal, consistent with literature precedent.[25, 26] Finally, withholding ADP but adding Mn^2+^ led to cell spreading but without platelet activation and this control also showed weak TaCT signal. Together, these experiments demonstrate that TaCT specifically tags platelets based on their molecular traction forces through integrin.

To showcase that TaCT is an effective method to report cell receptor forces, we used a Rho-associated protein kinase (ROCK) inhibitor, Y27632, to disrupt the force transmission through cytoskeleton and measure the dose-dependent response in MEF cells. Y27632 inhibits ROCK kinases by competing with ATP binding, and further result in decreased myosin activity, destabilization of actin filament and abolished stress formation (**Figure 1I**).[27] With Y27632 pre-treatment, MEF cells showed decreased Cy3B tension signal (**Figure 1J, Figure S7**). TaCT signal measured by flow cytometry showed a clear dose-dependent curve (**Figure 1K**) with an IC50= 773 nM, agreeing well with microscopy measurements and the literature reported IC50 values that range from hundreds of nM to low µM.[28]

After demonstrating TaCT with a single population of cells, we sought to demonstrate analysis of a binary mixture of cells with differing mechanical activity. We used parental MEF cells (wildtype, WT) and MEF cells with vinculin null (vin-) for this demonstration. Vinculin is an adaptor protein that localizes to FAs linking the cytoskeleton to adhesion receptors and aids in FA maturation (**Extended Figure 1A**).[29] We first performed TaCT on these two cell types separately. The vin-cells were stained with cell tracker dye CMFDA before plating onto TaCT substrates (**Figure 2A**). To ensure that the TaCT signal is not due to differential uptake, cholesterol uptake of WT and vin-cells was measured by flow cytometry in parallel with each TaCT assay (**Figure S8**). As shown in **Figure 2B** and **C**, WT cells had larger spreading area and produced more tension signal compared to vin-cells in the Cy3B channel. Flow cytometry also showed that there was more Atto647N signal in WT compared to vin-(**Figure 2D, Figure S8**) (WT: 26.6±3.7%, vin-4.5±0.3%). We then tested if this differential tension signal could be detected when the WT and vin-were co-cultured (**Figure 2E**). As expected, the WT cells displayed more TaCT tension signal by microscopy compared to vin-cells despite effectively spreading on the substrate (**Figure 2F**). The WT TaCT signal decreased slightly when these cells were co-cultured, with a 24.4±3.7% force positive population compared to 5.2±1.0% for vin-in the mixed population (**Figure 2G**). The minimal TaCT signal change observed for vin-cells in co-culture indicated that TaCT tags remain associated with target cells within the experimental time window (**Figure S8**). Together, these results confirm that TaCT can distinguish mechanically active subpopulations in heterogeneous mixtures of cells. To further demonstrate that TaCT produce a gradual response rather than a binary signal, we transfected MEF vin-cells with different amount of plasmid encoding GFP-vinculin (**Figure 2H**). We next investigated the relationship between vinculin expression and tension by quantifying GFP and Atto647 intensity at the single cell level. As expected, both the GFP and TaCT signal increased with increasing amount of the plasmid (**Figure S9**). Interestingly, the TaCT signal recovered with relatively low levels of GFP-vinculin, showing a linear regime followed by saturation of tension at increasing levels of vinculin, which indicates a complex relationship between focal adhesion tension and vinculin expression (**Figure 2I**).

**Figure 2.**
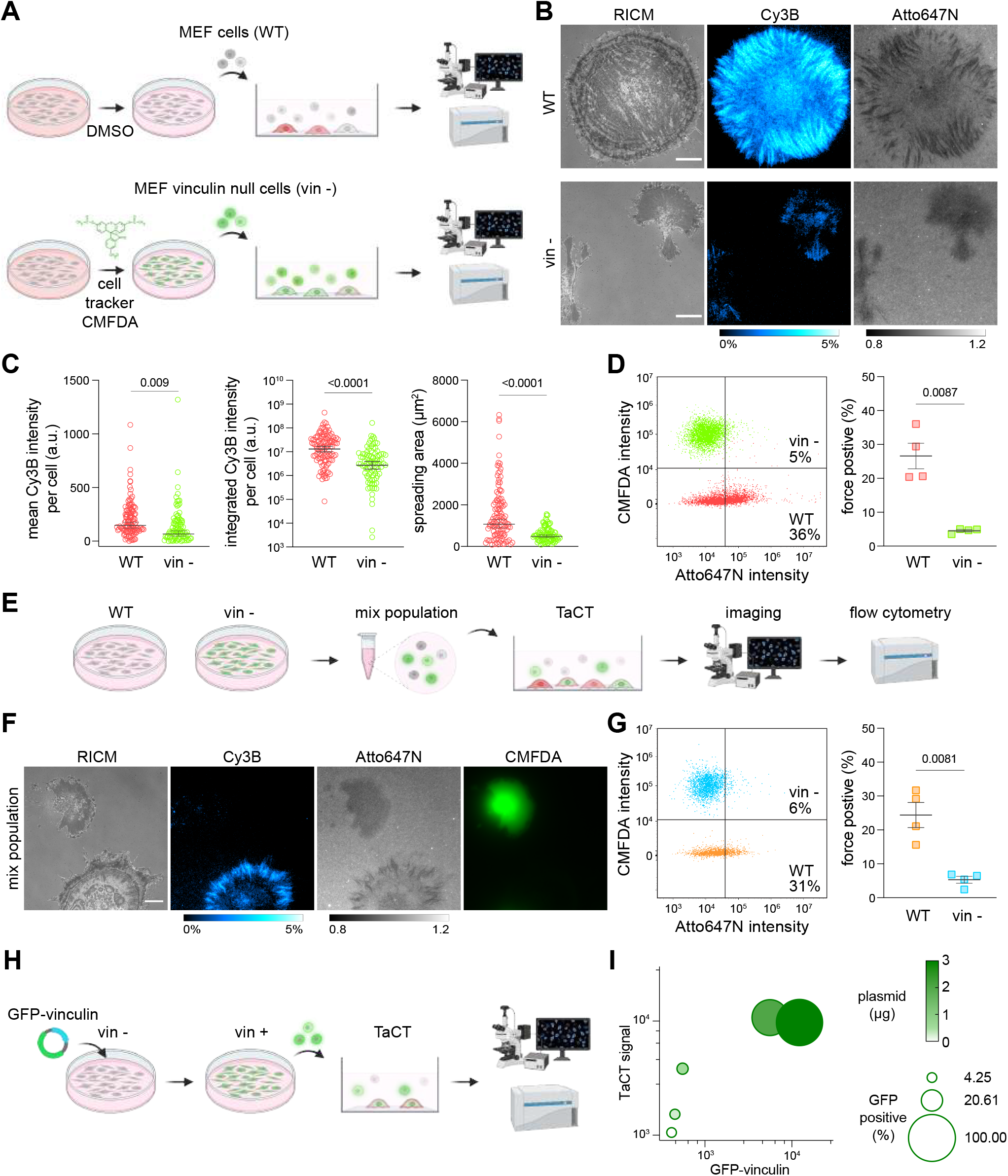
TaCT detecting mechanically active cells in a mixed population. (A) Schematic shows TaCT workflow for MEF WT and vin-cells. MEF vin-cells were stained with CMFDA for 15 min before plating onto TaCT substrate. TaCT probe microscopy imaging and flow cytometry analysis performed ∼1 h after cell seeding. (B) Representative RICM, Cy3B, and Atto647N microscopy images of WT and vin-cells seeded on TaCT substrates separately. Scale bar = 10 µm. (C) Plots of quantified microscopy images showed that tension signal (Cy3B, mean and integrated) and spreading area of WT cells were significantly higher than vin-cells (115 and 91 cells from n=3 replicates, two-tailed student’s t-test.) (D) Representative flow cytometry density plot and quantification of force positive cells show that the WT cells had significantly greater TaCT signal compared to vin-cells (n=4 replicates, mean ± SEM, two-tailed paired student’s t-test). (E) Schematic showing application of TaCT to co-cultured WT/vin-cells. (F) Representative RICM, Cy3B, and Atto647N microscopy images of co-cultured WT and vin-cells on TaCT substrate. Note that negative Atto647N signal for vin-cells was primarily due to optical effect from cell-surface contact. (G) Representative flow cytometry density plot and quantification of force positive cells show that the WT cells had significantly greater TaCT signal and force positive subpopulation compared to vin-cells incubated on the same TaCT substrate (n=4 replicates, mean±SEM, two-tailed paired student’s t-test). (H)Schematic describes the TaCT experiments using MEF vin-cells transfected with different amount of plasmid that encodes GFP-vinculin. (I) Plot shows the vinculin expression (x-axis, GFP MFI) versus TaCT signal (y-axis, Atto647N MFI) with different amount of plasmid transfected (n=4 replicates). Green color represents the amount of the GFP-vinculin plasmid, the size of the bubble represents the percentage of the GFP positive subpopulation.

In conclusion, we take advantage of 3’-5’ mediated DNA peeling to develop a new class of DNA tension probes to map the molecular forces generated by cells and to enable high-throughput flow-cytometry based detection of mechanically active cells. As is the case for all DNA tension probes, the probability of mechanical dehybridization is loading rate dependent. Therefore, to fully realize the potential of this technique, future investigations into the precise loading rate and force duration of mechanoreceptors is required and this needs to be coupled with the development of TaCT probes with different lengths and GC%. TaCT signal reports the total number of mechanical events exceeding *F*_peel_, which is orthogonal to biochemical analysis using antibodies and nucleic acids. If greater force magnitude detection is desired, duplexes with greater *F*_peel_ should be designed. In applications that use TaCT to characterize mechanically active cells, it is important to include a cholesterol strand uptake calibration to account for potential cholesterol insertion differences between different types of cells. A negative control that is not mechanically active should also be included to account for background uptake of cholesterol strands. Taken together, TaCT will open up a new class of tools for mechanobiology potentially allowing one to link single-cell mechanical phenotype with other biochemical properties.

**Extended Figure 1.**
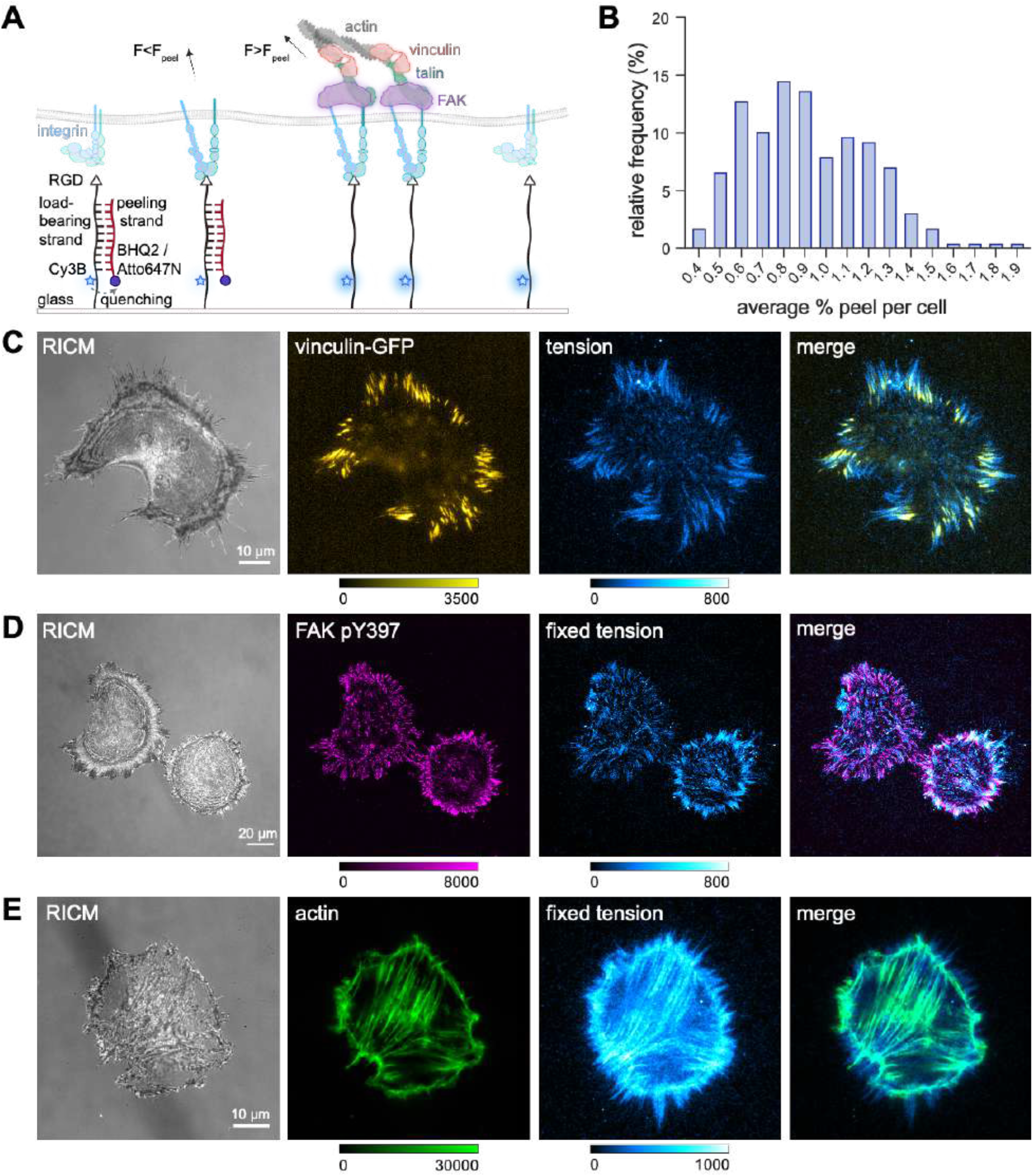
Mapping integrin forces with peeling probe. (A) Scheme showing that when integrin is in the inactive state, the peeling probe remains in the duplex form. As integrin binds and pulls on the RGD ligand, if F<F_peel_, the peeling strand is intact, and if F>F_peel_, the duplex dehybridizes and the Cy3B fluorescence turns on. FA proteins such as vinculin, talin, and FAK, as well as actin cytoskeleton participate intensively during the integrin force generation and mechanotransduction. (B) Histogram of 227 NIH3T3 cells showing the distribution of average % peel/µm^2^ per cell after 1 h incubation on peeling probe substrate. Data acquired from 3 biological replicates, bin width = 0.1%. (C) Representative microscopy images show that GFP-vinculin colocalized with integrin tension signal. Images were acquired with MEF GFP-vinculin cells that were cultured on peeling probe substrate. (D) Representative microscopy images show that the phosphorylated FAK (pY397) colocalized with fixed integrin tension signal. MEF cells were incubated on peeling probe substrate for 40-45min, fixed and stained with Rabbit anti-FAK pY397 and Alexa488 labeled secondary antibody, followed by imaging. (E) Representative microscopy images show that the actin stress fibers colocalized with fixed tension signal. MEF cells were incubated on peeling probe substrate for 60-90 min, fixed and stained with Alexa647-phalloidin, followed by imaging.

**Extended Figure 2.**
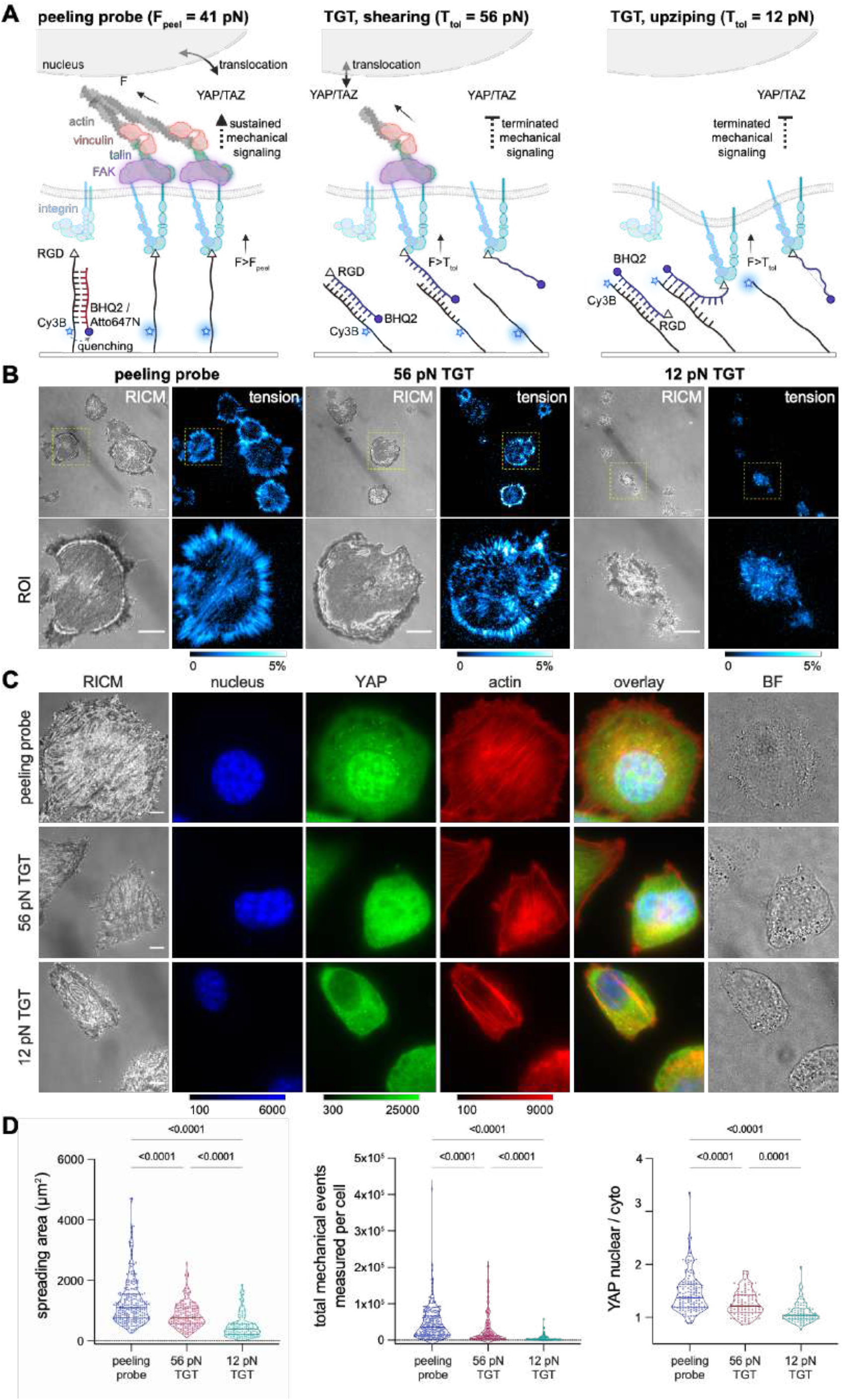
Peeling probe does not perturbate mechanotransduction unlike TGTs. (A) Schematics comparing the mechanism of TaCT/peeling probe and TGTs. Peeling probe maps integrin tension greater than 41 pN as the BHQ2 strand is separated, and the RGD anchor remains despite the duplex dehybridization. In contrast, TGTs map integrin forces greater than 56 or 12 pN in the shearing or unzipping geometry. The force-induced rupture of the duplex generates a Cy3B turn-on fluorescence signal as the top BHQ2 strand separates from the Cy3B anchor strand. Unlike peeling strand, the loss of the RGD anchors in TGTs terminates mechanotransduction through integrins. (B) Representative microcopy images of NIH3T3 cells incubated on peeling probe or TGT substrates for ∼45-60 min. Second row of images show the zoom-in view of the ROIs marked with the yellow dashed box. (C) Microscopy images of NIH3T3 cells incubated on peeling probe or TGT substrates for ∼90 min, fixed and stained for nucleus (DAPI), actin (SirActin), and YAP (Alexa488 conjugated antibody). Scale bar = 5 µm. (D) Quantitative analysis of the spreading area, %peel or %rupture, and YAP translocation for NIH3T3 cells incubated on three substrates for 60 min. For spreading area and tension quantification, data was collected from 3 biological replicates (n = 227, 150, and 105 for cells on peeling probe, 56 pN TGT, and 12 pN TGT substrate). For analysis on YAP translocation, images in DAPI channel were used as masks to quantify the mean fluorescence intensity of nuclear YAP and cytoplasm YAP. Data was acquired from 3 biological replicates (total n = 134, 98, and 86 cells for peeling probe, 56 pN TGT and 12 pN TGT substrate). Plots show lines at the median and interquartile values. Statistical analysis was performed with one-way ANOVA and Tukey’s multiple comparison test.

**Extended Figure 3.**
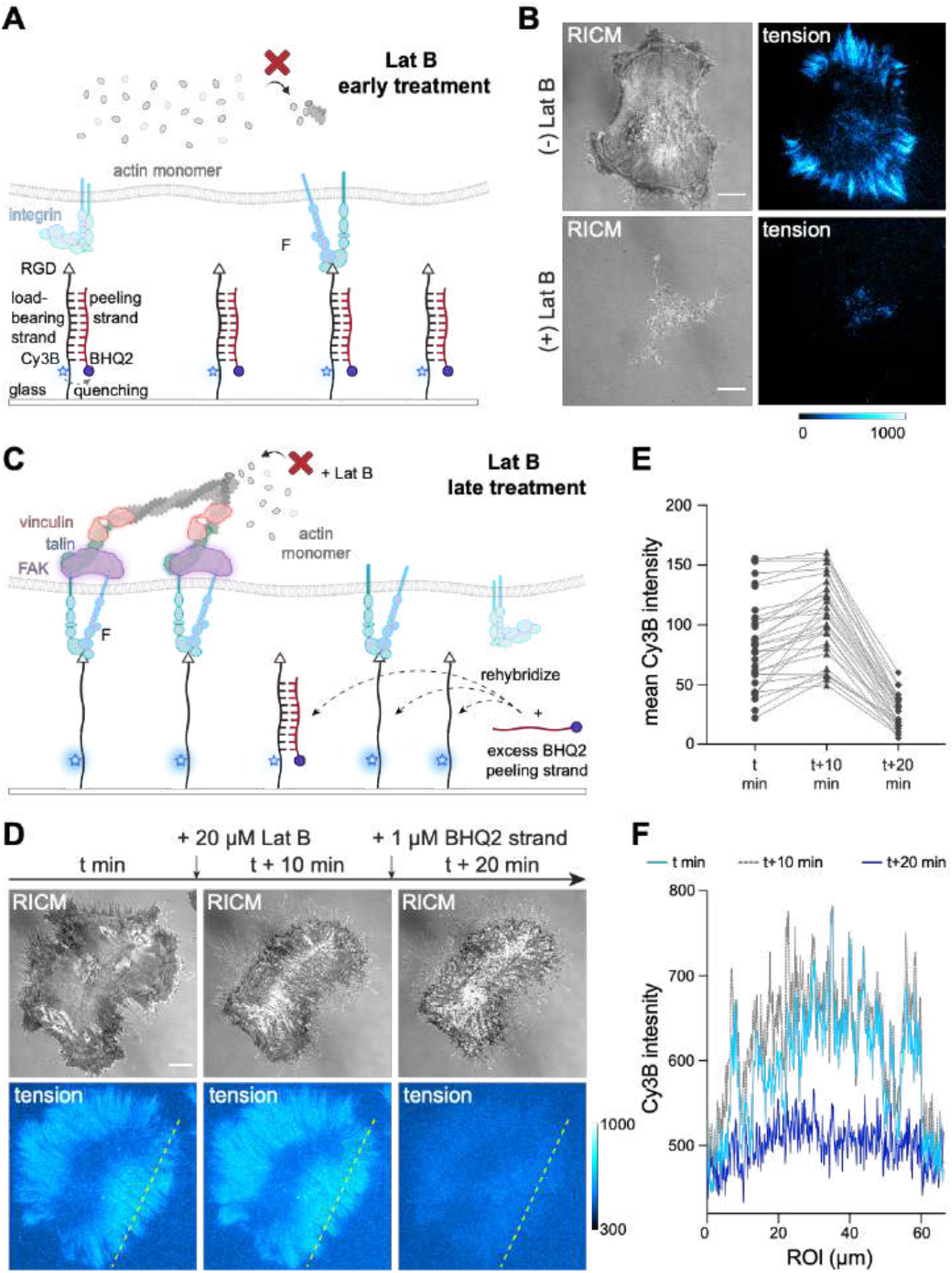
Lat B inhibition of cells. (A) Scheme showing that Lat B early treatment inhibits integrin force generation by inhibiting actin polymerization. (B) Representative RICM and Cy3B microscopy images of MEF cells incubated on peeling probe substrate after early Lat B treatment. Cells were treated with 20 µM Lat B after 15 min of plating on the substrate and imaged after 45 min of incubation. (C) Scheme showing that when integrin force signals are already generated on peeling probe substrate, if the force transmission is terminated by late Lat B treatment, with additional BHQ2 peeling strand the peeling probe can rehybridize to the duplex form. (D) Representative RICM and raw Cy3B microscopy images of MEF cells treated with 20 µM Lat B 50 min after seeding. The tension signal remained after 10 min of Lat B treatment and diminished after the addition of excess BHQ2 peeling strand. (E) Quantitative analysis of tension signal changes after Lat B treatment and the addition of BHQ2 peeling strand in n = 30 cells. (F) Linescan of the ROI (yellow dashed line) in (D) before Lat B treatment, with Lat B treatment, and with excess BHQ2 peeling strand.

**Extended Figure 4.**
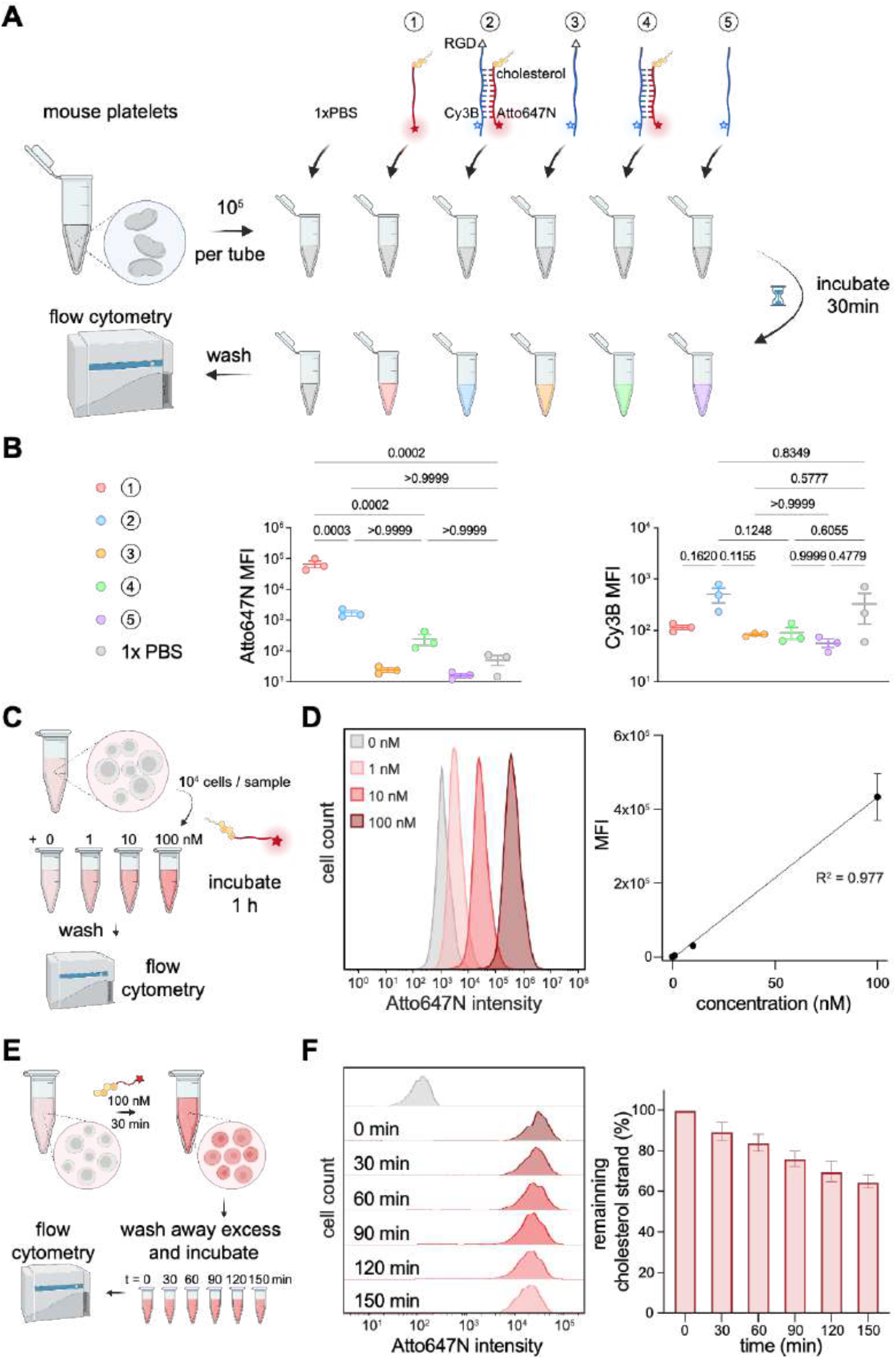
Cholesterol DNA strands association and dissociation in cells. (A) Scheme showing flow cytometry measurements of the DNA strands uptake in solution. Mouse platelets were incubated with 50 nM of cholesterol peeling strand, TaCT duplex, load-bearing strand, TaCT duplex lacking RGD, and load-bearing strand lacking RGD in Tyrode’s buffer for 30 min and spun down three times to wash away the excess oligos. The association was measured in both Atto647N and Cy3B channels by a flow cytometer. (B) Cy3B and Atto647N median fluorescence intensity (MFI) of platelets incubated with different oligonucleotides. Data collected from 3 replicates (mean ± SEM). Statistical analysis was performed by one-way ANOVA and Tukey’s multiple comparison. (C) Scheme showing flow cytometry measurements of the concentration dependent incorporation of cholesterol peeling strand. MEF cells were incubated with 0, 1, 10, and 100 nM of cholesterol peeling strand for 1 h. The excess cholesterol peeling strand was washed away by spinning down in PBS three times, and the fluorescence intensity of cells was measured by a flow cytometer. (D) Representative histogram of cholesterol peeling strand association in cells. Atto647N MFI was plotted from 3 replicates (mean±SD). Linear relationship between cholesterol strand concentration and cell association was found, R^2^ = 0.977. (E) Scheme showing the measurement of cholesterol peeling strand dissociation from the cell. NIH3T3 cells were incubated with 100 nM cholesterol peeling strand for 30 min and rinsed with PBS 3 times. Cells were divided into 6 aliquots and the remaining cholesterol strand in cells was measured every 30 min for 150 min by flow cytometry. (F) Representative histogram shows the decay of fluorescence in cells over time. The normalized MFI from 2 replicates was plotted to show the dissociation of cholesterol strand over time in NIH3T3 cells (mean±SD).

**Extended Figure 5.**
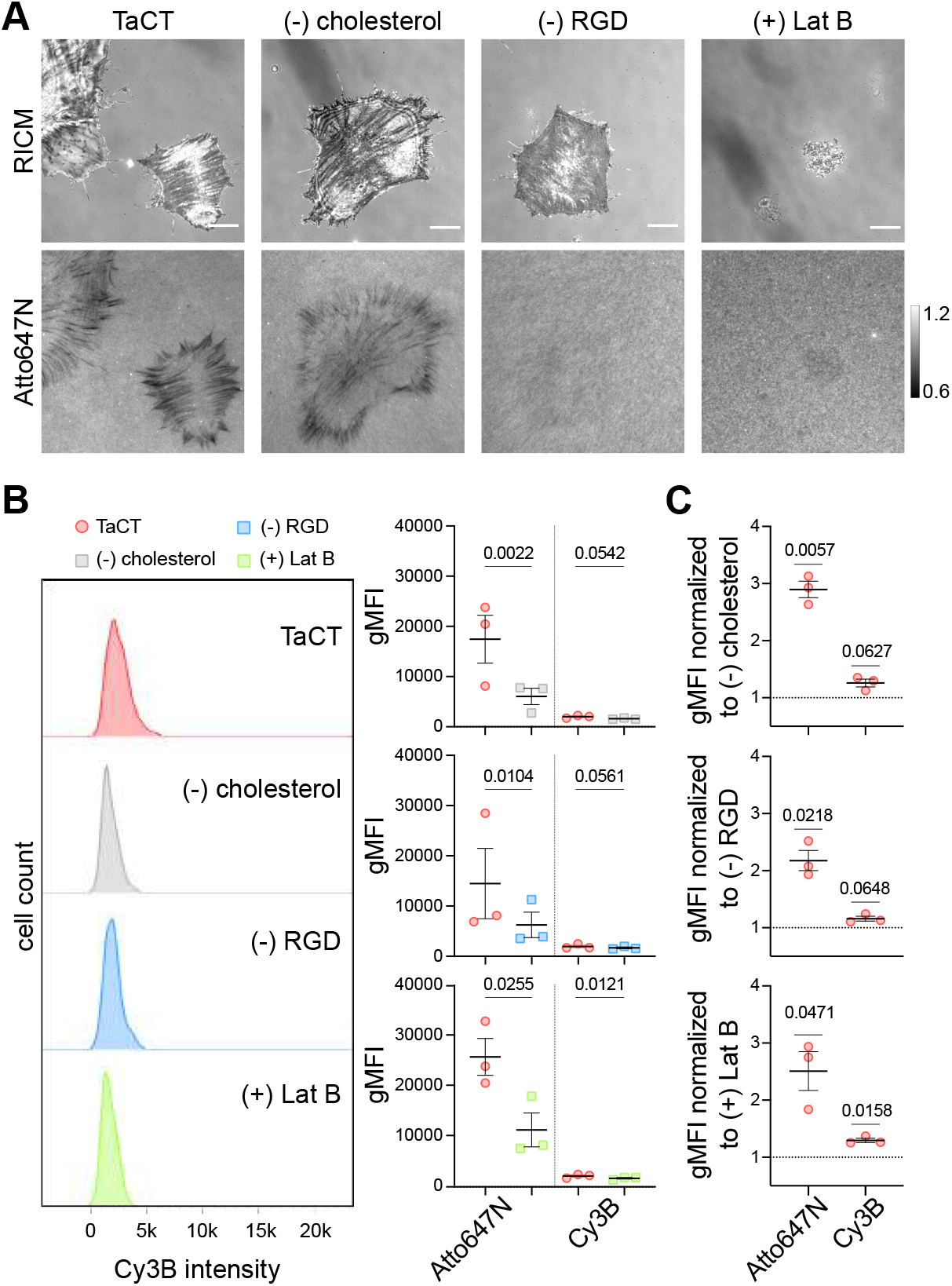
Control experiments supporting the concept of TaCT. (A) Representative microscopy images of NIH3T3 cells cultured on TaCT substrate, or control substrate lacking cholesterol, lacking RGD, or treated with Lat B. Atto647N shows the signal normalized to the background intact probes. Scale bar = 10 µm. (B) Representative flow cytometry histograms show that NIH3T3 cells in TaCT, (-)cholesterol, (-)RGD, and (+)Lat B groups had indistinguishable Cy3B intensity. Quantitative analysis (mean±SEM) shows that cells cultured on TaCT substrate had similar level of geometric mean fluorescence intensity (gMFI) compared to control groups in Cy3B channel, despite higher Atto647N gMFI. Data collected from 3 replicates for each control, and statistical analysis was performed using ratio paired two-tailed student’s t-test. (C) Plots show the gMFI of cells after TaCT normalized to different controls in both Atto647N and Cy3B channels (n = 3, mean±SEM). One sample t-test was used for statistical analysis to test whether TaCT signal is significantly different in Atto647N and Cy3B compared to a hypothetical value of 1 (representing no TaCT signal in the corresponding control group). TaCT signal in Atto647N channel consistently provides 2 to 3 fold higher gMFI.

**Extended Figure 6.**
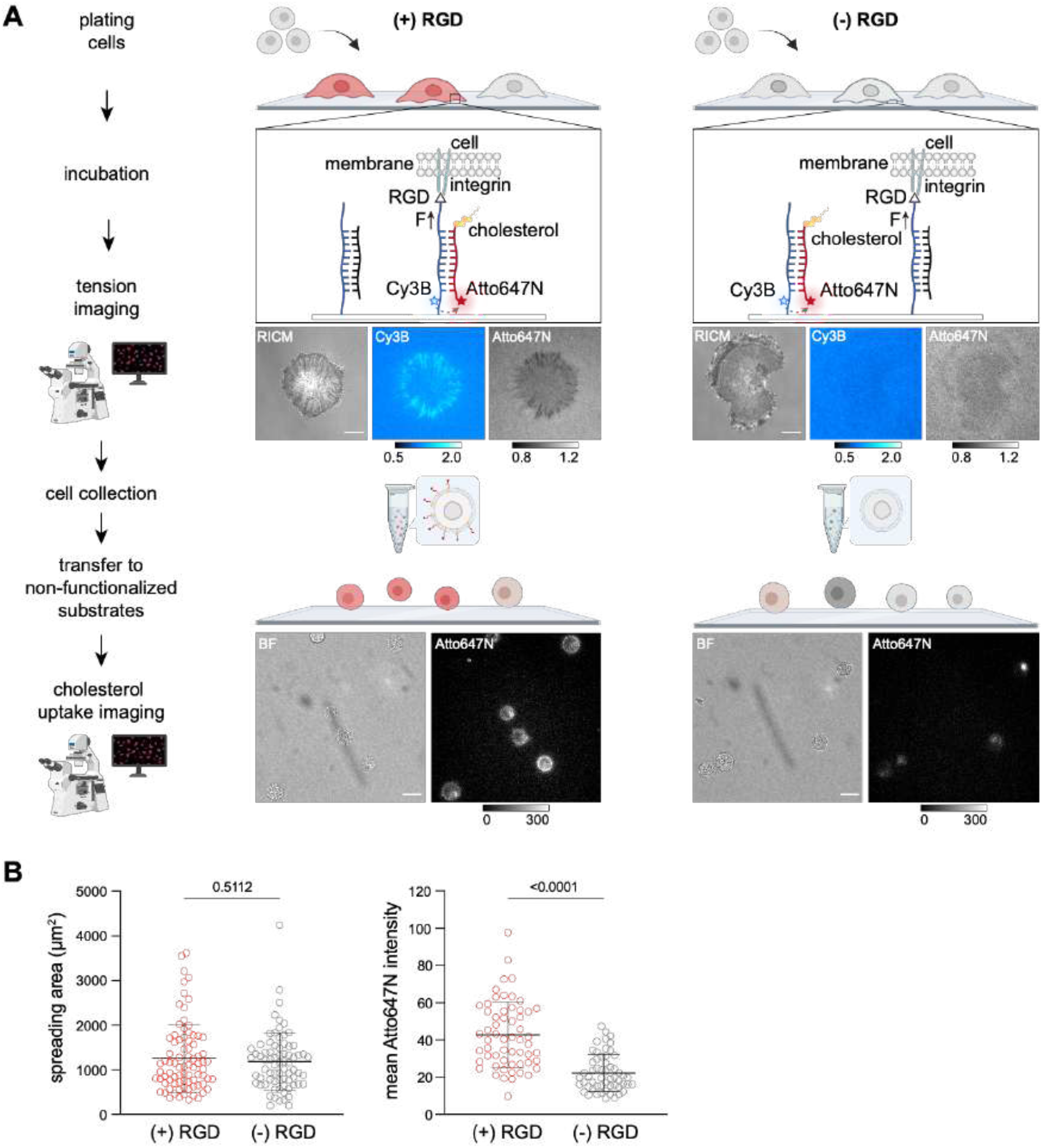
Spreading control for TaCT. (A) Scheme and representative microscopy data showing that TaCT signal is not due to spreading of the cells on the substrate. NIH3T3 cells were incubated on a TaCT substrate doped with a non-fluorescent DNA duplex, or a TaCT substrate lacking RGD doped with a non-fluorescent DNA duplex presenting the RGD motif, and imaged 45 min - 60 min after seeding. After confirming the cell spreading, cells were collected, rinsed, and added to non-fluorescent glass substrate to image the cholesterol peeling strand incorporation in the cells. Tension images were normalized to the background of intact probes. Scale bar = 10 µm. (B) Quantitative analysis of the cell spreading area (n=82 and 74 cells) and cholesterol peeling strand incorporation (n=58 and 56 cells) for cells incubated on TaCT (+)RGD or (-)RGD substrates. Data was collected from three replicates and statistical analysis was performed using two-tailed student’s t-test.

**Extended Figure 7.**
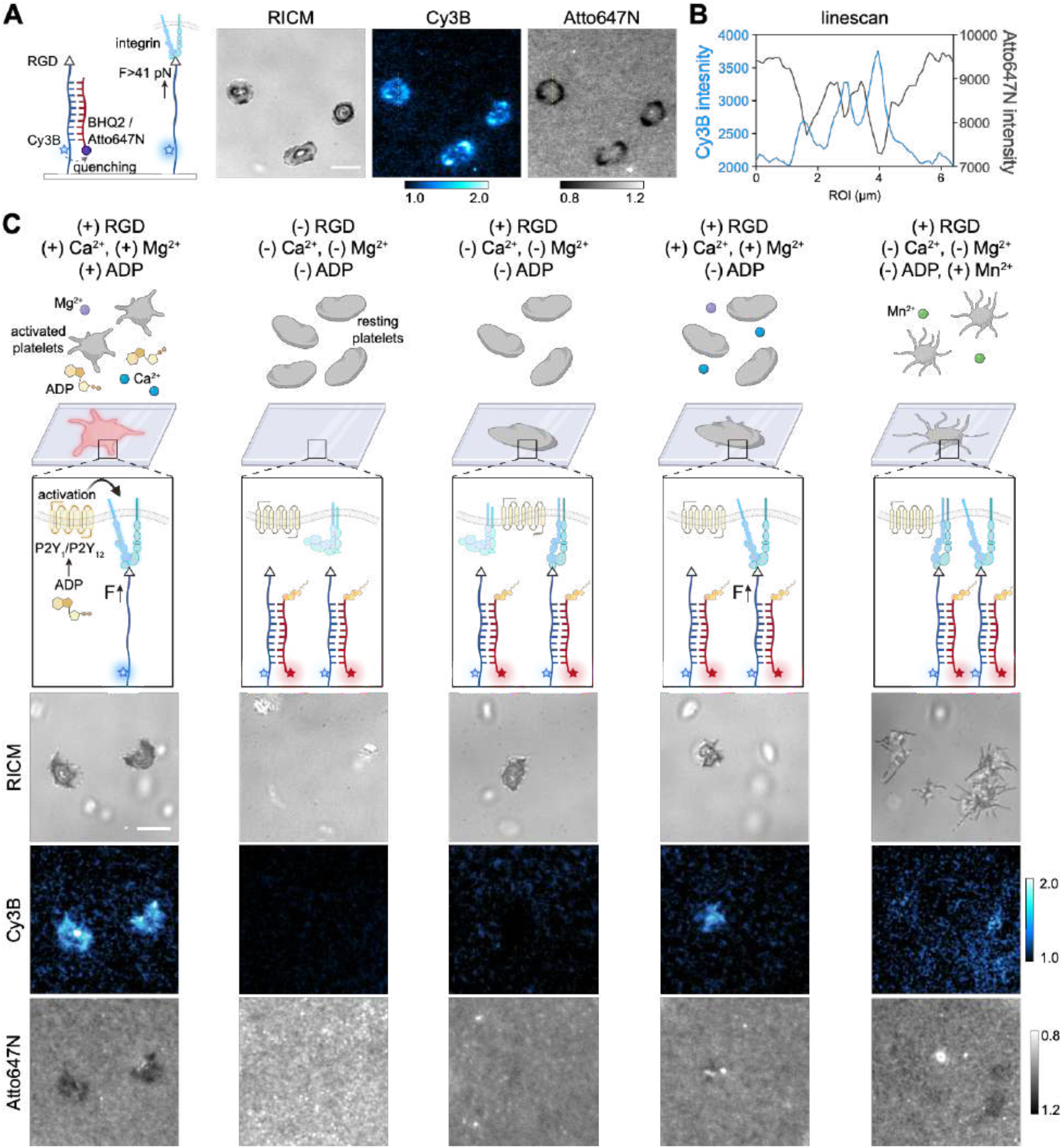
TaCT in Platelets. (A) Scheme and representative microscopy images showing tension mapping with peeling probe in mouse platelets. Tension images were normalized to the background of intact peeling probes, scale bar = 5 µm. (B) Linescan of the ROI (yellow dashed line) shows anti-colocalization of Cy3B and Atto647N intensities from raw data. (C) Scheme and representative images show mouse platelets incubated on the TaCT substrate in different buffer conditions. RGD, Ca^2+^, Mg^2+^, and ADP leads to platelets activation, and withholding any of them results in no/impaired activation. Tension images were normalized to the background of intact TaCT probes, scale bar = 5 µm.

## Data availability statement

Data summary file contains all individual replicate data from main text, all individual replicate data from supporting information, and all p values from statistical analyses performed are provided with figures in this manuscript.

## Code availability statement

Code used for oxDNA data analysis and graph generation is publicly available online at https://github.com/SalaitaLab/Tension_activated_cell_tagging

## Author contributions

R.M. and K.S. conceived the idea and wrote the manuscript. R.M., A.V., A.R., and B.R.D. carried out experiments and data analysis. A.V. performed oxDNA simulation. W.C., R.L., and B.P. provide mouse platelets. All authors contributed to editing the manuscript.

## Supporting Information

**Table S1.** Representative list of methods for mechanophenotyping. The majority of past reports describing approaches for mechanical phenotyping use deformability as a marker. Cell deformation is induced by a variety of methods, including narrow constrictions, laser traps, shear stress, and acoustic waves, etc. The table below lists representative papers using deformability as well as adhesion strength or traction forces-based mechanophenotyping methods. Note that all past methods are based on cell-response rather than molecular tension generated by specific adhesion receptors.

| Marker | Article Title | Year | Journal |
| --- | --- | --- | --- |
| Deformability | Optical Deformability as an Inherent Cell Marker for Testing Malignant Transformation and Metastatic Competence[1] | 2005 | *Biophysical Journal* |
|  | Analyzing cell mechanics in hematologic diseases with microfluidic biophysical flow cytometry[2] | 2008 | *Lab on a chip* |
|  | Microfluidics-Based Assessment of Cell Deformability[3] | 2012 | *Anal. Chem.* |
|  | Microfluidic micropipette aspiration for measuring the deformability of single cells[4] | 2012 | *Lab on a chip* |
|  | Characterizing deformability and surface friction of cancer cells[5] | 2013 | *PNAS* |
|  | Pinched-flow hydrodynamic stretching of single-cells[6] | 2013 | *Lab on a chip* |
|  | Quantitative Diagnosis of Malignant Pleural Effusions by Single-Cell Mechanophenotyping[7] | 2013 | *Science Translational Medicine* |
|  | Real-time deformability cytometry: on-the-fly cell mechanical phenotyping[8] | 2015 | *Nature Methods* |
|  | Microfluidic cell sorting by stiffness to examine heterogenic responses of cancer cells to chemotherapy[9] | 2018 | *Cell Death & Disease* |
|  | High-throughput microfluidic compressibility cytometry using multi-tilted-angle surface acoustic wave[10] | 2021 | *Lab on a chip* |
|  | ... etc[11-13] |  |  |
| Cell adhesion strength | Probing Cell Adhesion Profiles with a Microscale Adhesive Choice Assay[14] | 2017 | *Biophysical Journal* |
|  | High-Throughput Characterization of Cell Adhesion Strength Using Long-Channel Constriction-Based Microfluidics[15] | 2021 | *ACS sensors* |
| Cell traction force | High-throughput screening for modulators of cellular contractile force[16] | 2015 | *Integrative Biology* |
|  | Elastomeric sensor surfaces for high-throughput single-cell force cytometry[17] | 2018 | *Nature Biomedical Engineering* |

**Supplementary Note. Peeling probe offers significant advantages as a tension sensor compared to TGTs**.

Unlike TGTs, the force-induced denaturation of the peeling probe does not result in termination of the mechanical force experienced by adhesion receptors (**Extended Figure 2A**). Note however that TGTs and peeling probes are chemically similar as they are primarily comprised of DNA duplexes. We compared cell spreading area, integrin tension maps (**Extended Figure 2B)**, and the mechanical signaling outcomes for cells incubated on peeling probe or TGT substrates (**Extended Figure 2C**). Quantitative analysis (**Extended Figure 2D**) showed that the spreading area for cells incubated on the peeling probe substrate was significantly larger than that of cells incubated on 12 and 56 pN TGT substrates. The total number of mechanical events was calculated by multiplying the probe density, cell area, and the %peel or %rupture of cells incubated on the three different substrates. There were significantly more total mechanical pulling events for cells incubated on peeling probe substrate versus TGTs, possibly due to two reasons: first, the *F*_peel_ is ∼41 pN, which is less than the T_tol_ for 56 pN TGT; and second, loss of integrin-ligands for TGT substrates modulates cell signaling. Based on the differences in the cell spreading area and tension profile, we sought to further analyze mechanical signaling by quantifying Yes-associated protein (YAP) translocation to the nucleus. YAP translocation to the nucleus is regulated by focal adhesion signaling, F actin organization, and mechanotransduction. YAP nuclear signaling is critical in regulating cell morphology, proliferation, and plasticity.[18] YAP staining showed that cells incubated on peeling probe substrates had the highest YAP nuclear/cyto ratio, followed by 56 pN and then 12 pN TGTs. This result confirms that terminating mechanotransduction through integrin dampens YAP translocation into the cell nucleus, which impairs transcription activation.[19] This result suggests that the peeling probe is better suited to decouple tension sensing from receptor force manipulation within cells in mechanobiology studies, which is highly advantageous.

## 1. Materials

### 1.1 Oligonucleotides

The oligonucleotides used in this study are listed in **Table S2**.

**Table S2.**
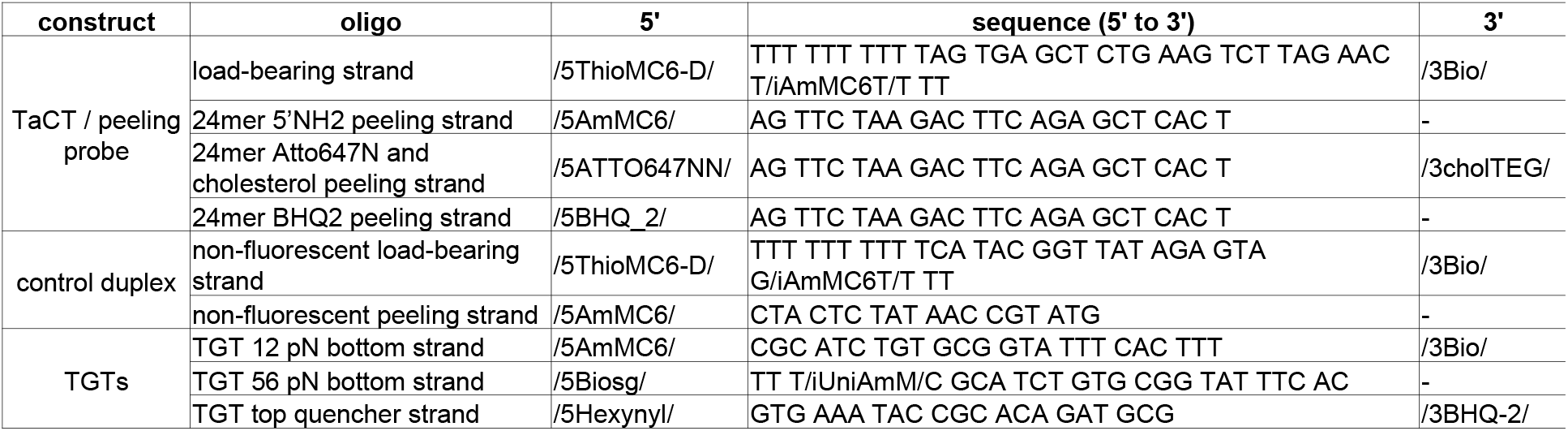
List of oligonucleotides.

### 1.2 Reagents

The reagents used in this study are listed in **Table S3**.

**Table S3.**
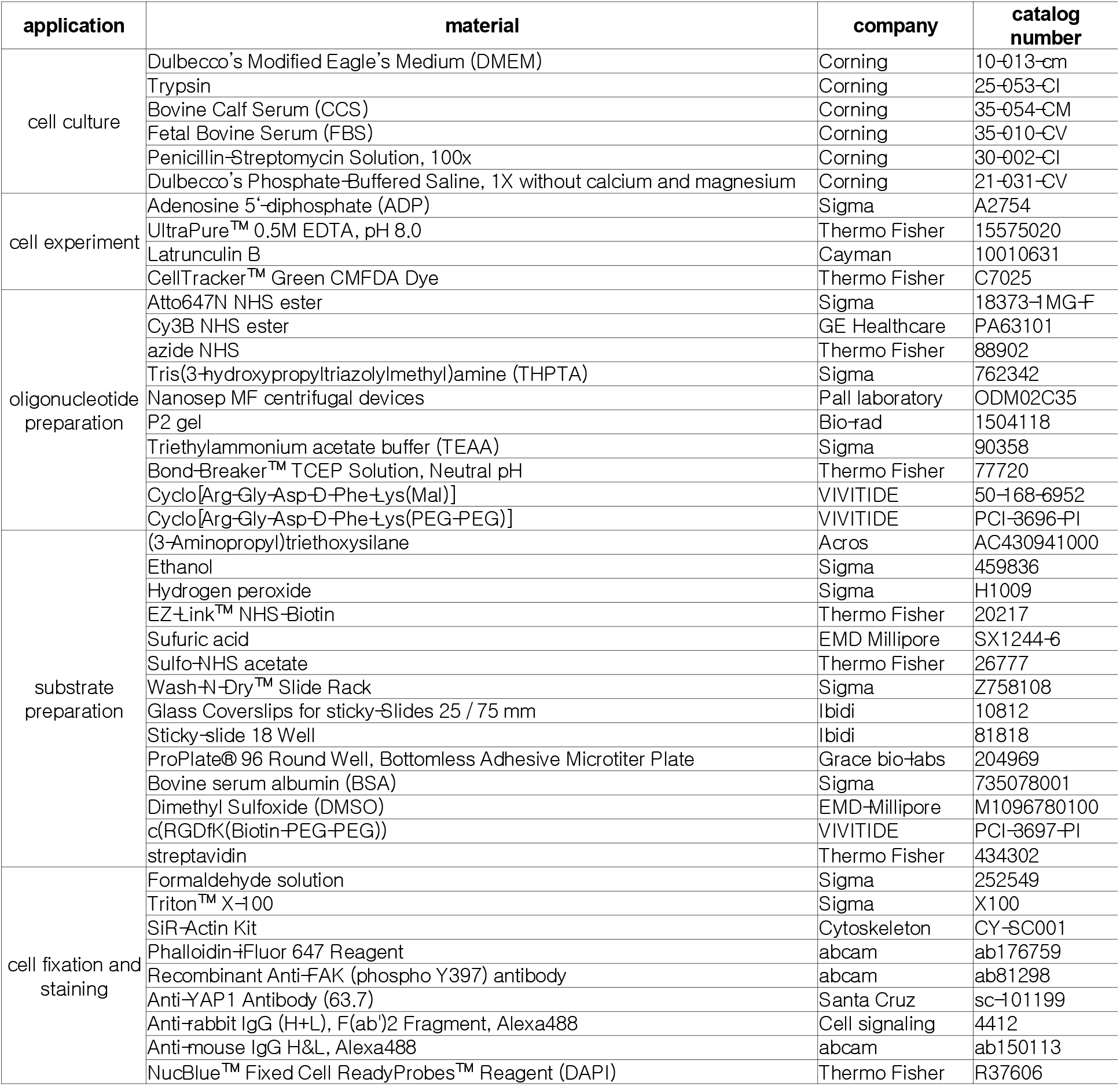
List of reagents.

### 1.3 Equipment

The equipment used in this study is listed in **Table S4**.

**Table S4.** List of equipment.

| equipment | company |
| --- | --- |
| Barnstead Nanopure water purifying system | Thermo Fisher |
| Centrifuges | Eppendorf |
| AdvanceBio Oligonucleotide C18 column, 4.6 x 150 mm, 2.7 μm | Agilent |
| Grace Alltech C18 column | Grace |
| High-performance liquid chromatography, 1100 | Agilent |
| Nanodrop 2000 UV-Vis Spectrophotometer | Thermo Fisher |
| CFI60 Apochromat TIRF 100X Oil Immersion Objective Lens, N.A. 1.49 | Nikon |
| Prime 95B-25mm Back-illuminated sCMOS Camera. 1608x1608,30fps | Photometrics |
| Nikon Ti2-E Motorized Research Microscope | Nikon |
| Ti2-ND-P Perfect Focus System 4 | Nikon |
| SOLA SE II 365 Light Engine | Nikon |
| NIS Elements software | Nikon |
| C-FL Surface Reflection Interference Contrast (SRIC) Cube, Excitation:535/80nm (495-575nm), Dichroic Mirror: 400nm, No Emission Filter | CHROMA |
| CF-L AT CY5/Alexa Fluor 647/Draq 5 Filter Set, Excitation: 620/50nm(595-645nm), Emission: 690/50nm (665-715nm), Dichroic Mirror: 655nm | CHROMA |
| C-FL DS Red Hard Coat, High Signal-to-Noise, Zero Shift Filter Set,Excitation: 545/30nm (530-560nm), Emission: 620/60nm (590-630nm),Dichroic Mirror: 570nm | CHROMA |
| CF-L AT EGFP/FITC/Cy2/Alexa Fluor 488 Filter Set, Excitation: 480/30nm(465-495nm), Emission: 535/20nm (525-545nm). Dichroic Mirror: 505nm | CHROMA |
| CytoFLEX V0-B3-R1 Flow Cytometer (equipped with a 488 and a 638 nm laser) | Beckman Coulter |

## 2. Methods

### 2.1 Simulation of DNA peeling

The dsDNA peeling was modeled with oxDNA by adding harmonic traps (effectively springs) to two terminal nucleotides of a DNA stand (**Figure S1**). Each trap was assigned with a stiffness constant of 11.42 pN/nm, and one of the traps was moved at a given rate (loading rate = 2.81×10^3^ nm/s) with respect to the other fixed trap. The effective stiffness constant of the two traps in series can be calculated using:

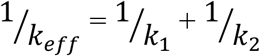

where *k*_1_ and *k*_2_ are the stiffness constants of the two traps and *k*_*eff*_ is the effective stiffness constant. The *k*_*eff*_ of the system is calculated to be 5.71 pN/nm. The total displacement of the two terminal nucleotides from the respective trap centers is defined as net displacement. Force is then calculated by multiplying *k*_*eff*_ with net displacement projected along the force axis (z -axis). This force is plotted against the net displacement with an exponential moving average (EMA) of data points. The oxDNA model considers base pairs as broken when the pairing energy is less than 10% of that of a fully formed base pair. The critical force at which peeling occurs (*F*_peel_) is defined here as the force at which the total number of base pairs falls to ≤5. The first 100 data points were averaged (to reduce the stochasticity in simulation data) to obtain the value of *F*_peel_. Modeling parameters were adopted from published literature and from examples available at dna.physics.ox.ac.uk website.[20]

### 2.2 Oligonucleotide preparation

#### Conjugation with dyes

The load-bearing strand was conjugated with Cy3B NHS on the internal amine, the 24mer peeling strand was conjugated with Atto647N NHS on the 5’ amine, and the TGT bottom strands were conjugated with Cy3B (**Table S2**). Briefly, an excess amount of NHS dye (50 µg) was dissolved in DMSO immediately before use and then reacted with 10 nmol amine oligonucleotide at room temperature for 1 h in 1×PBS containing 0.1 M NaHCO_3_. The mixture after reaction was desalted with P2 gel and purified with an HPLC coupled to an Advanced oligonucleotide C18 column (**Figure S2**). The product was eluted in Solvents A: 0.1 M TEAA and B: acetonitrile with linear gradients of 10-35% Solvent B over 25 min and 35-100% Solvent B over 5 min at 0.5 mL/min flow rate, and dried in a Vacufuge overnight. The dried oligo-dye conjugate was reconstituted in water and the concentration was determined by its absorbance at 260 nm with Nanodrop.

#### Conjugation with cRGD

Maleimide-cRGDfk was used for conjugation with thiol oligonucleotide strand (load-bearing strand). Briefly, 5 nmol of thiol oligonucleotide was reduced in 200× molar excess TCEP at room temperature for 15 min, and the mixture was added to 0.5 mg (excess) of maleimide-cRGDfk in 1×PBS (pH = 6.8) to react at room temperature for 1 h. The reaction mixture was then desalted with P2 gel and purified by HPLC with advanced oligonucleotide column as described above.

For alkyne oligonucleotide TGT top quencher strand, azide-RGD was first prepared and then conjugated to the oligo. Briefly, excess amount of azide-NHS (∼0.5 mg) was used to react with 100 µg cyclic (RGD)fk-PEG2-amine overnight at 4 °C. The product was purified with HPLC coupled to a Grace C18 column for peptide purification. Product was eluted in Solvents A: 0.1% TFA in H_2_O and B: 0.1% TFA in ACN with linear gradients of 10-40% Solvent B over 30 min and 40-75% Solvent B over 10 min at 1 mL/min flow rate. The purified product was dried in a Vacufuge overnight and characterized with Maldi-TOF-MS (data not shown). Stock solutions of CuSO_4_ (20 mM in water), THPTA [Tris(3-hybroxypropyltriazolyl methyl)amine] (50 mM in water), and sodium ascorbate (100 mM in water) were prepared. A final mixture of 100 µM azide-RGD, 50 µM of alkyne-DNA, 0.1 mM CuSO_4_, 0.5 mM THPTA, and 5 mM sodium ascorbate in 1X PBS was allowed to react at room temperature for 2 h. The product was purified by P2 gel, followed by HPLC with advance oligonucleotide column as described above.

### 2.3 DNA substrates preparation

#### Amine glass slides

Glass slides (25 × 75 mm) were placed on a Wash-N-Dry rack, rinsed with water (18.2 MΩ), and sonicated in ethanol and water for 15 min each, followed by 6 rinses with water. Fresh piranha solution was made by mixing concentrated sulfuric acid and hydrogen peroxide (30%) at a 3:1 ratio (v/v) in a total volume of 200 mL and added to the slides for etching. CAUTION: PIRANHA SOLUTION IS HIGHLY REACTIVE AND MAY EXPLODE IF MIXED WITH ORGANIC SOLVENTS. Next, the slides were rinsed again with copious amount of water to remove the acid, and then rinsed in ethanol to remove water. Then, 200 mL of 3% APTES in ethanol was prepared and added to the glass slides at room temperature to react for 1 h for amine modification. After reaction, the glass slides were washed with copious amounts of ethanol, and bake dried in an oven (80 °C) for 20 min. The amine modified glass slides were stored at -20 °C until use.

#### Biotin substrate preparation

An amine modified glass slide was carefully placed on a parafilm-lined petri dish. Then, 1 mL of 4 mg/mL Biotin-NHS in DMSO was added to the slide and incubated overnight. On the second day, after washing with copious amounts of ethanol and then water several times, the glass slide was air-dried, and attached to an ibidi sticky-slide imaging chamber or a bottomless adhesive 96-well plate. The wells were passivated in 1% BSA in PBS for 10 min at room temperature and then washed with PBS. Streptavidin at 50 µg/mL in PBS was added to each well and incubated for 30 min at room temperature, and the excess was washed away with PBS. Meanwhile, TaCT/peeling probe (load-bearing strand:peeling strand = 1:1.5) was annealed at 50 nM by heating to 95 °C for 5 min and then gradually cooling down to 20 °C in 20 min. For TGT substrates, the BHQ2 top strand was annealed with Cy3B bottom strand (1.1:1 ratio) at 50 nM. Next, the probes were added to each streptavidin-coated well and incubated for 30 min. The excess was rinsed away with PBS before imaging (**Figure S3**).

### 2.4 Probe density calibration

#### Lipid vesicles

Lipid vesicles were prepared with 100% DOPC or with 99.5% DOPC and 0.5% Texas Red DHPE (TR-DHPE). Lipid was dissolved in chloroform in a round-bottom flask and dried with rotary evaporator and under ultra-high purity N_2_ stream. The dried lipids were subsequently resuspended in Nanopure water at 2 mg/mL and endured three freeze/thaw cycles in acetone and dry ice bath or 40 °C water bath. Lipids were then extruded ten times through a high-pressure extruder assembled with a 0.2 μm membrane to obtain uniform vesicles.

#### Supported lipid bilayer preparation

A glass bottom 96-well plate was used for preparing supported lipid bilayers. Each well was filled and soaked with ethanol for 15 min and rinsed with 5 mL Nanopure water. Subsequently, 200 µL of 6 M NaOH solution was added to each well for base etching at room temperature for 1 h. After washing each well with 10 mL Nanopure water, lipid mixtures containing different percentage of TR-DHPE vesicles (0.01%-0.5%) were added at 0.5 mg/mL in PBS to coat the glass surfaces for 20 min. After the lipid vesicles fused to the surfaces, the wells were rinsed with 10 mL water and 10 mL PBS and imaged with a fluorescence microscope to create a standard curve for probe density calibration.

#### Solution phase standard curve preparation

Surfaces were first washed with ethanol and water and passivated with 1% BSA to prevent any surface adsorption. Different concentrations of TR-DHPE (2.96 nM – 740 nM) and a series of Cy3B-oligo solutions (10 nM – 500 nM) were then added to each well and their bulk fluorescence intensity in solution was measured with a fluorescence microscope (4 µm above surface) to create a standard curve (**Figure S4**).

### 2.5 Cell culture and transfection

NIH3T3 cells were cultured in DMEM (10% CCS, 1% P/S) at 37 °C with 5% CO_2_. MEF and MEF vinculin null cells were cultured in DMEM (10% FBS, 1% P/S) at 37 °C with 5% CO_2._ Cells were passaged at 80% confluency every two days by detaching using trypsin and replating at lower density. For MEF vinculin null cells, a plasmid encoding full-length vinculin and GFP was used for transfection. Cells were seeded in a 6 well plate and transfected with 0, 0.25, 0.5, 2, or 3 µg of the plasmid for 24 h. The expression was validated by flow cytometry (data not shown).

### 2.6 Mouse platelets

Blood of C57Bl/6J mice were collected after cardiac puncture or via retro-orbital plexus, and anticoagulated with acid-citrate-dextrose. After mixing with equal volume of Tyrode’s buffer (140 mM NaCl, 2.7 mM KCl, 0.4 mM NaH_2_PO_4_, 10 mM NaHCO_3_, 5 mM Dextrose, 10 mM HEPES, 3U apyrase), mouse platelets were isolated by centrifugation at 200×g for 5 min and subsequently supplemented with 1U apyrase and 1 µM prostaglandin E1. Finally, the mouse platelets were spun down at 700×g for 5 min and resuspended in Walsh buffer (137 mM NaCl, 2.7 mM KCl, 1 mM MgCl_2_, 3.3 mM NaH_2_PO_4_, 20 mM HEPES, 0.1% glucose, 0.1% bovine serum albumin, pH 7.4) at a density of ∼1×10^9^ platelets per mL.

### 2.7 Fluorescence microscopy

#### Integrin tension imaging

Imaging was conducted with a Nikon Ti2-E microscope. All the fibroblast cells were plated onto the DNA probe substrates in DMEM (10% serum, 1% P/S) and allowed to attach for 15-20 min in the incubator at 37 °C in medium. Then, the cells were allowed to further spread at room temperature and imaged up until 60 min after plating in RICM, Cy3B, and Atto647N channels with accommodating filter settings and a sCMOS detector.

Mouse platelets were plated onto the DNA probe substrates in 1× Tyrode’s buffer (134 mM NaCl, 12 mM NaHCO_3_, 2.9 mM KCl, 0.34 mM NaH_2_PO_4_, 5 mM glucose, 5 mM HEPES, 0.1% BSA, pH 7.4) supplemented with 2 mM CaCl_2_, 1 mM MgCl_2_, and 10 µM ADP, and imaged at room temperature up until 60 min after plating in RICM, Cy3B, and Atto647N channels.

#### Immunostaining

Cells incubated on DNA probe substrates were fixed in 4% formaldehyde for 10 min, followed by two gentle PBS washes and permeabilization with 0.1% triton X-100 in PBS for 10 min. The cells were then blocked in 1% BSA for 1 h and then stained. Fixed cells were incubated with primary antibodies (anti-YAP: 2 µg/mL, anti-pFAK Y397: 5 µg/mL in 1% BSA) at 4 °C overnight. After rinsing three times with PBS, cells were incubated in secondary antibody (2 µg/mL in 1% BSA) for 1 h at room temperature. For actin straining, fixed cells were incubated with 0.5 µM SiR-actin or phalloidin iFluor 647 (1:1000 dilution). Microscopy images were acquired with accommodating filter settings.

### 2.8 Flow cytometry

For all fibroblast cells, 1×10^4^ cells were seeded on the TaCT substrate and collected by gentle scraping 60 min after seeding. After spinning down and washing in PBS containing 5 mM EDTA, the collected cells were resuspended in DPBS (without Ca^2+^ and Mg^2+^, with 5 mM EDTA) and immediately run through a flow cytometer for TaCT analysis.

For mouse platelets, 3 µL of the platelets were diluted in 600 µL 1× Tyrode’s buffer. 100 µL platelets were then added to the TaCT substrate with different supplements (Ca^2+^, Mg^2+^, ADP, Mn^2+^). After 1 h, platelets were collected by pipetting them off from the substrate and then washed before running through flow cytometer for TaCT analysis.

### 2.9 Data analysis Tension

#### imaging data

All microscopy data was analyzed with Image J software.

For quantification of FRET efficiency, Cy3B load-bearing strand and Atto647N peeling strand were annealed and immobilized on a biotin substrate and imaged to obtain the fluorescence intensity with both donor and acceptor present (I_DA_). Similarly, Cy3B load-bearing strand and amine peeling strand were annealed and immobilized on a biotin substrate and imaged to obtain fluorescence intensity when only the donor was present (I_D_). After sCMOS background subtraction of the images, the fluorescence intensity in Cy3B was averaged from 5 different positions of the substrates. The FRET efficiency was calculated with the equation 1 – I_DA_/I_D_.

The probe density was estimated as described previously in literature[21]. Briefly, peeling probe and TGT substrates with unquenched Cy3B fluorescence strand were prepared. A calibration curve was made with fluorescent supported lipid bilayers (SLB) to provide a count for molecules/µm^2^, and the F-factor was calculated using a standard curve with a series of Cy3B concentrations (**Figure S4**). The average intensity of three substrates was used to calculate the probe density using the calibration curve.

Microscopy images of integrin tension was presented in formats including raw data, background subtracted or normalized molecular tension images, and %peel/%rupture. For quantitative analysis of microscopy data with cells, the sCMOS background was subtracted, and the fluorescence intensity (mean±SD) of the substrate background was used as a threshold. Raw integrated intensity, contact area, and tension area of the ROIs of the cells were measured and plotted. The tension signal in Cy3B for each image was used to calculate the %peel or %rupture according to literature (**Figure S10**) [21].

#### Flow cytometry data

Flow cytometry data was analyzed with Flowjo software. Briefly, the debris and aggregated cells were gated out by first identifying the live cells using the forward scatter and side scatter area and then the singlet cells using the forward and side scatter height (**Figure S10**). Following gating, the fluorescence signal of each viable singlet cell in the Atto647N channel and Cy3B channel was measured and analyzed. The signal intensity was presented in histograms, and the median fluorescence intensity (MFI) or geometric mean fluorescence intensity (gMFI) was used for comparison between groups. The %positive cells was determined by creating a vertical gate in the histogram of the negative control group so that ∼99.5% of cells in the negative control had a lower fluorescence than the gate.

### 3. Figures

**Figure S1.**
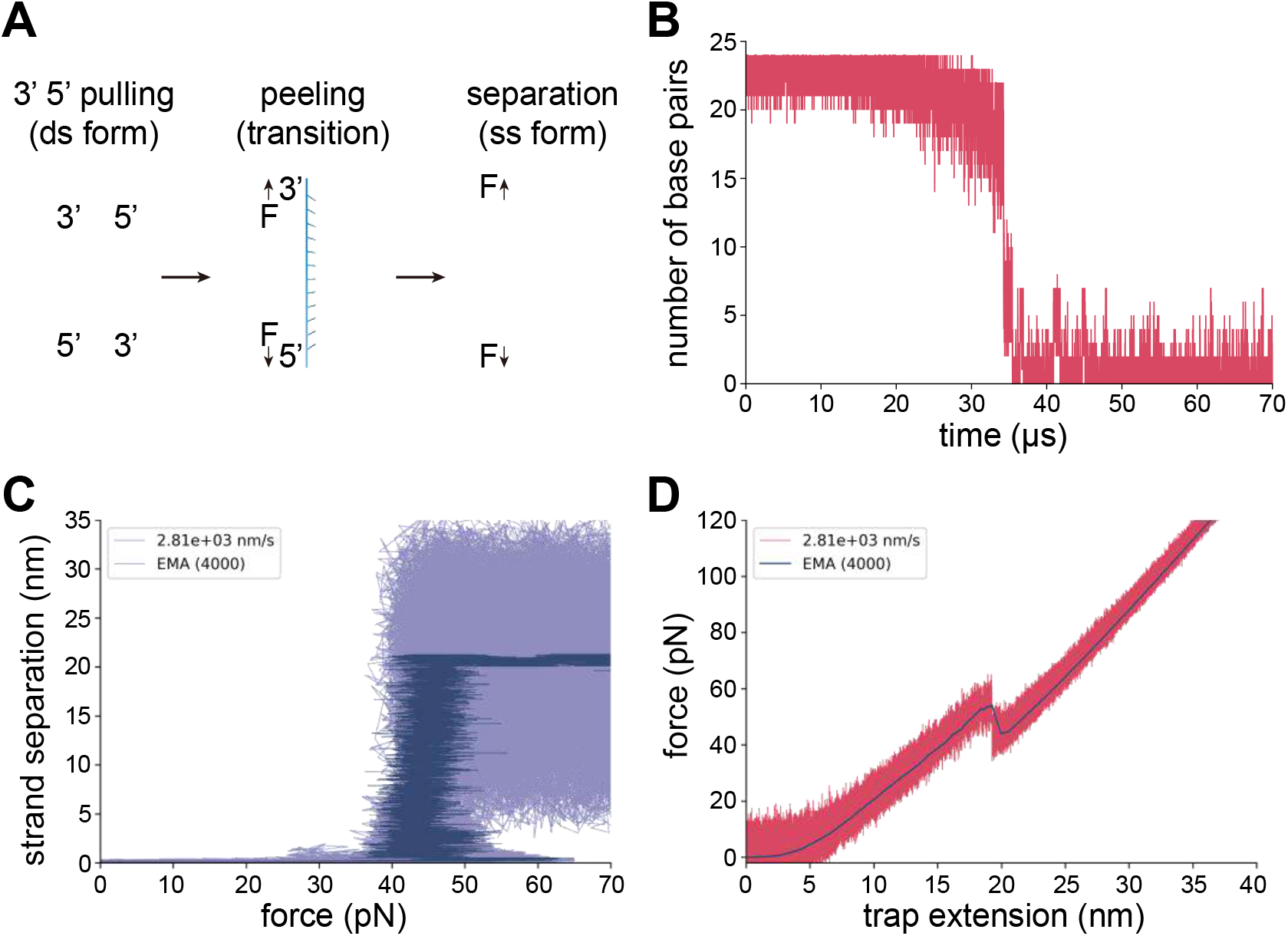
Simulations with oxDNA. (A) Scheme showing the force-induced peeling mechanism of oligonucleotides. (B) Simulation data of the transition state during force-induced peeling. The transition occurs on µs timescale, which further indicates that the transition state of peeling probe can be ignored while using it as a digital tension sensor. (C) Simulation result of the strand separation distance versus force. (D) Simulated force-distance curve. Conditions used for simulation: loading rate = 2.81×10^3^ nm/s, ionic strength = 0.156 M Na^+^, and effective stiffness constant k_eff_ = 5.71 pN/nm. The details of the simulation are further described in the methods section.

**Figure S2.**
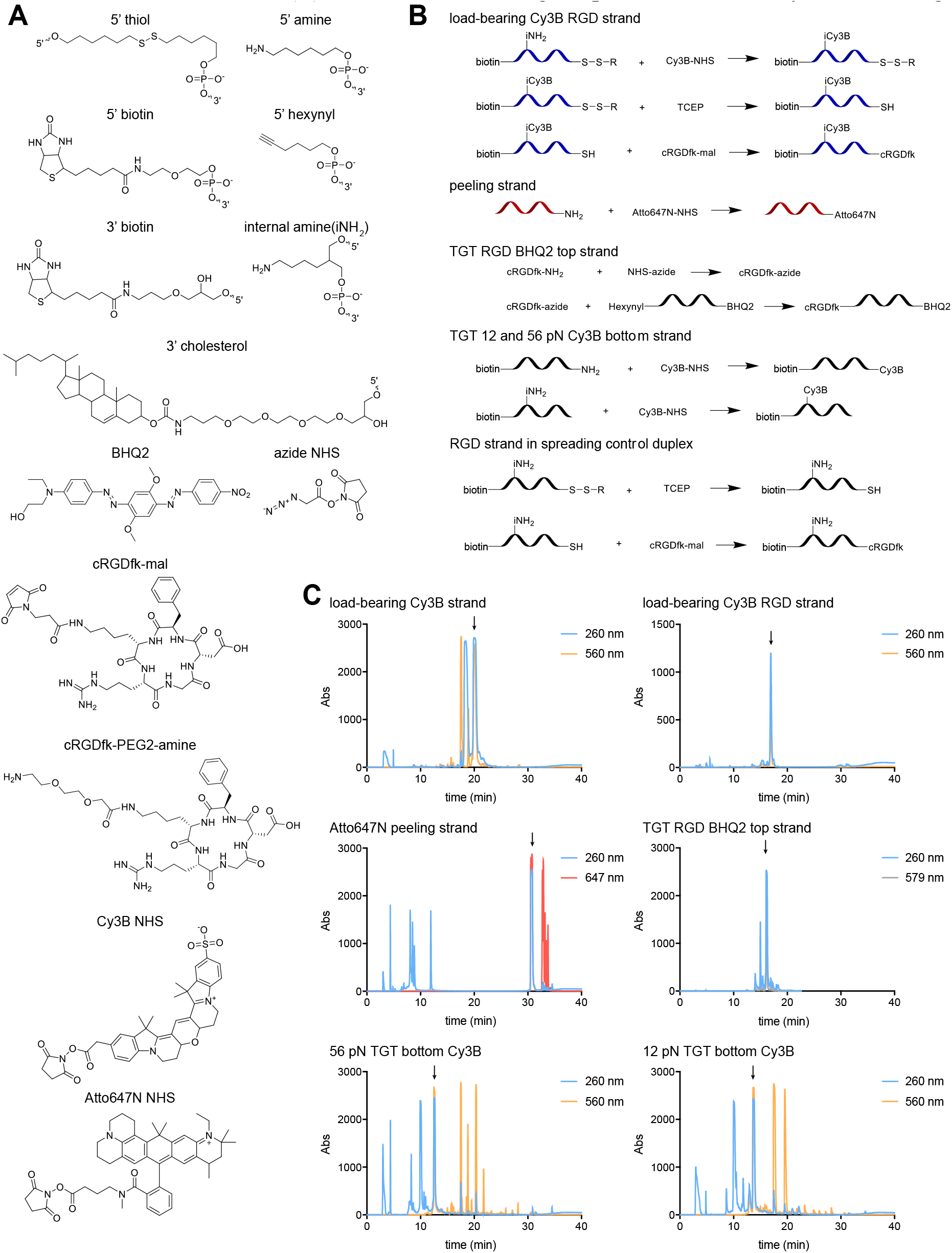
Oligonucleotide preparation. Chemical structures of the modifications on the oligonucleotides. (B) Schemes showing the modification of DNA strands. (C) HPLC traces showing the purification of the synthesized oligos.

**Figure S3.**
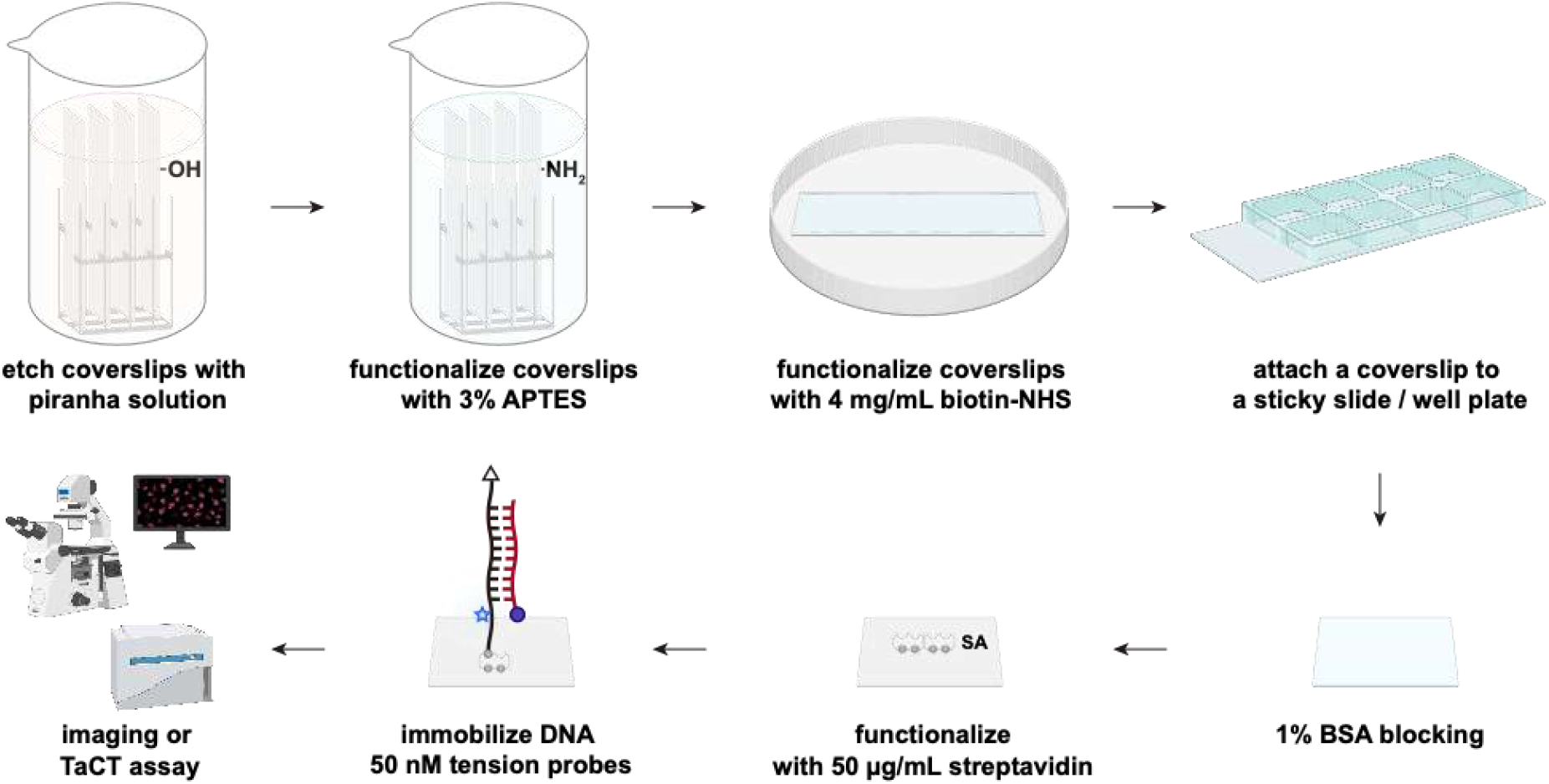
Surface functionalization. Glass slides were etched in piranha solution and functionalized with 3% APTES. Biotin-NHS was used to react with the amines on the glass slides. After attaching biotinylated glass slide to a sticky slide or a sticky well plate, surfaces were blocked with 1% BSA. Streptavidin was added to bind to the biotin on the surface, and the DNA probes were attached to the surfaces via biotin-streptavidin interaction. After surface preparation, the tension probe substrates were used for tension imaging or TaCT assay.

**Figure S4.**
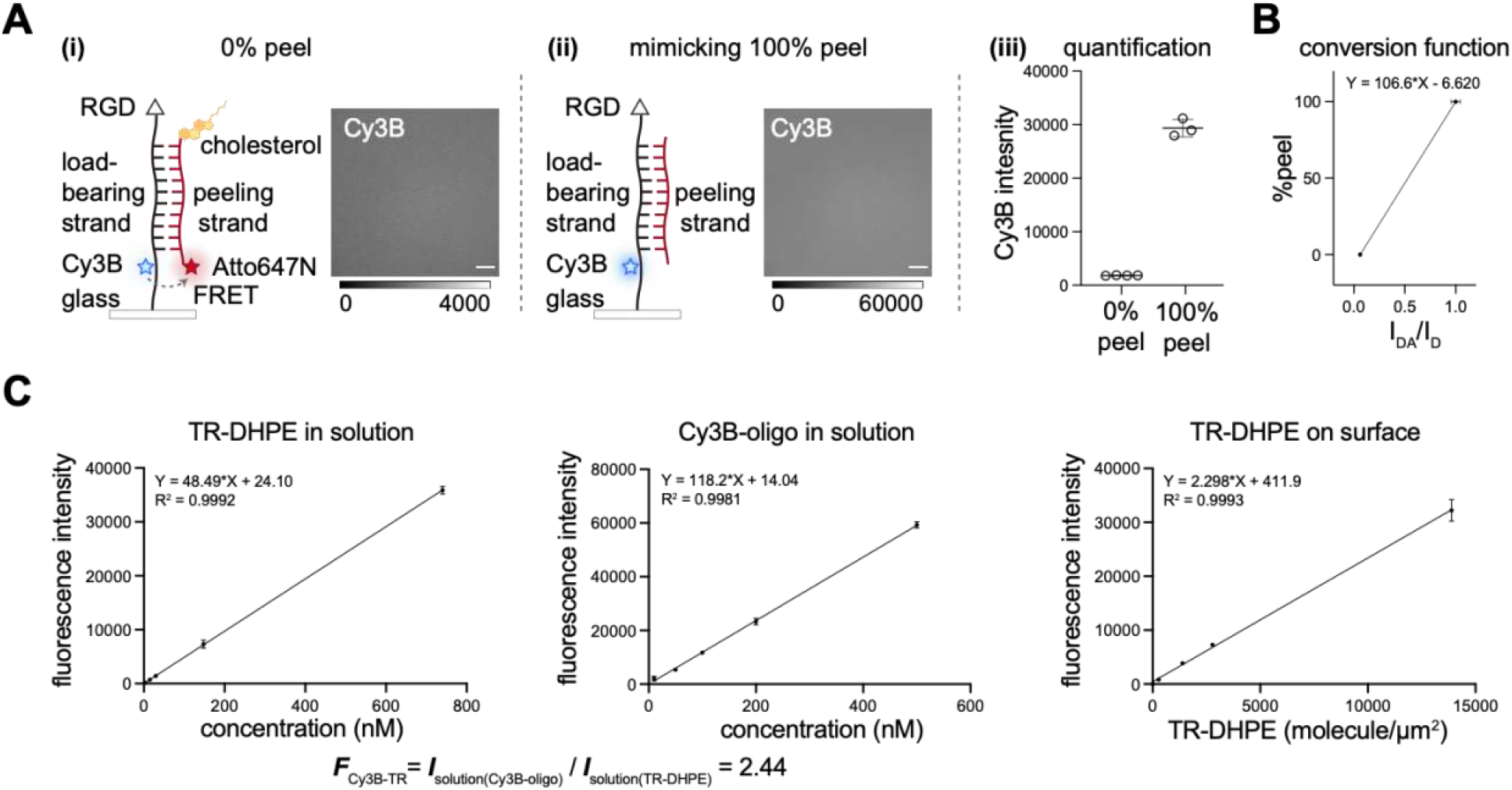
TaCT substrate characterization. (A) Scheme and representative microscopy images of the TaCT substrate (i) with the acceptor Atto647N (0% peel, I_DA_), or (ii) without the acceptor Atto647N (mimicking 100% peel, I_D_). Scale bar = 20 µm. The Cy3B fluorescence intensity of the substrate (iii) was quantified from multiple XY coordinates from > 3 independent replicates (mean±SD). The FRET efficiency was calculated to be 93.8% using the equation 1 – I_DA_/I_D_, where I_DA_ is the fluorescence intensity with the donor Cy3B and the acceptor Atto647N present, and I_D_ is the fluorescence intensity with the donor Cy3B only. (B) The conversion between I_DA_/I_D_ and %peel. Briefly, I_DA_, the fluorescence intensity of the 0% peel substrate (i), and I_D_, the fluorescence intensity of 100% peel substrate (ii) were used to generate the conversion function to find the %peel from the fluorescence tension signal. (C) Probe density quantification on TaCT substrates. Lipid standards were prepared by mixing DOPC and TR-DHPE at different percentages, and the fluorescence intensity was measured within the linear range for TR-DHPE in solution and TR-DHPE lipid bilayers on surface to create standard curves. A series of Cy3B-oligo solutions at different concentrations was prepared and their fluorescence intensity in solution was measured to create a standard curve. Data was plotted from 3 replicates (mean±SD). The scaling factor ***F***_Cy3B-TR_ was calculated using the equation ***F***_Cy3B-TR_ = I_solution(Cy3B-oligo)_/I_solution(TR-DHPE)_, where I is the concentration normalized fluorescence intensity of the standards (slope).[22] The TR-DHPE molecules per µm^2^ was estimated assuming the footprint of a lipid is 0.72 nm^2^ and was used for creating the standard curve to find I_surface(TR-DHPE)_ (note the factor 2 due to the nature of lipid membrane)[23], and the scaling factor ***F***_Cy3B-TR_ was applied to find the normalized fluorescence intensity I_surface(Cy3B-oligo)_ of the Cy3B-oligo on surface. The probe density was estimated to be 5242±286 probes/µm^2^ from the Cy3B fluorescence intensity of the substrates functionalized with duplexes lacking Atto647N (**Figure S4A, ii**).

**Figure S5.**
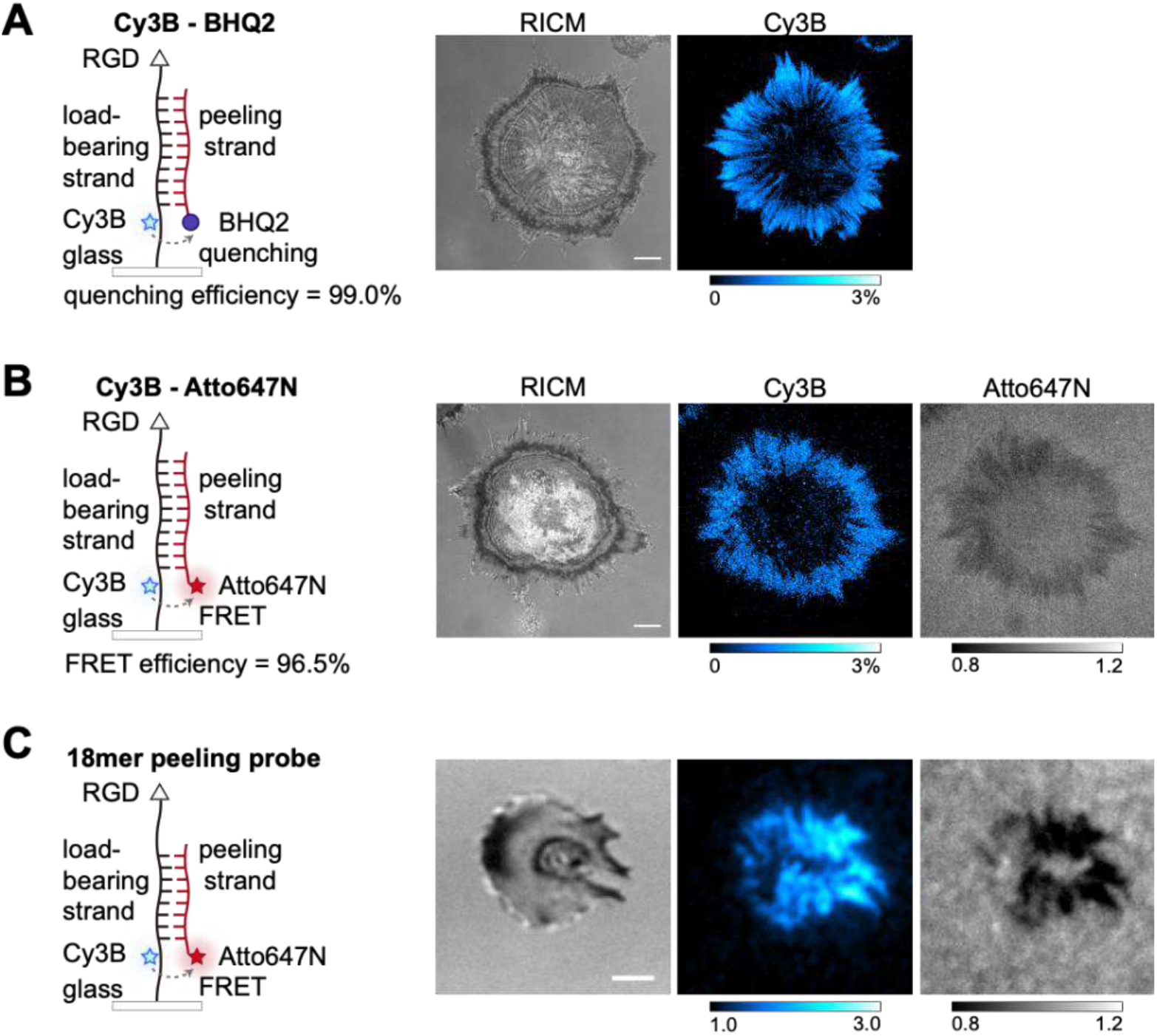
Iterations of DNA probes based on peeling mechanism for microscopy studies on integrin tension. (A) Scheme shows the tension probe design with only BHQ2 on the peeling strand. Representative microscopy images in RICM and Cy3B channel show integrin tension produced by MEF cells ∼45 min after plating. Scale bar = 10 µm. (B) Scheme shows the tension probe with only Atto647N on the peeling strand. Representative microscopy images in RICM, Cy3B, and Atto647N channel show integrin tension produced by MEF cells ∼45 min after plating. Scale bar = 10 µm. Quenching efficiency and FRET efficiency for iterations in (A) and (B) were calculated using the same method described in **Figure S4** and were used for microscopy data processing. (C) Scheme shows the tension probe with a 18mer duplex region and with only Atto647N on the peeling strand. Representative microscopy images in RICM, Cy3B, and Atto647N channel show integrin tension produced by mouse platelets ∼45 min after seeding. Scale bar = 2 µm. Tension signal was normalized to background fluorescence intensity. We included representative images of the 18mer peeling probe to demonstrate the modularity of the probe. This class of DNA probes is a general design to study molecular tension transmitted through surface receptors. One can change the ligands or change surface chemistry for different cell types to accommodate the need for receptor tension mapping, and presumably one can tune the oligonucleotide to achieve differential *F*_peel_.[24]

**Figure S6.**
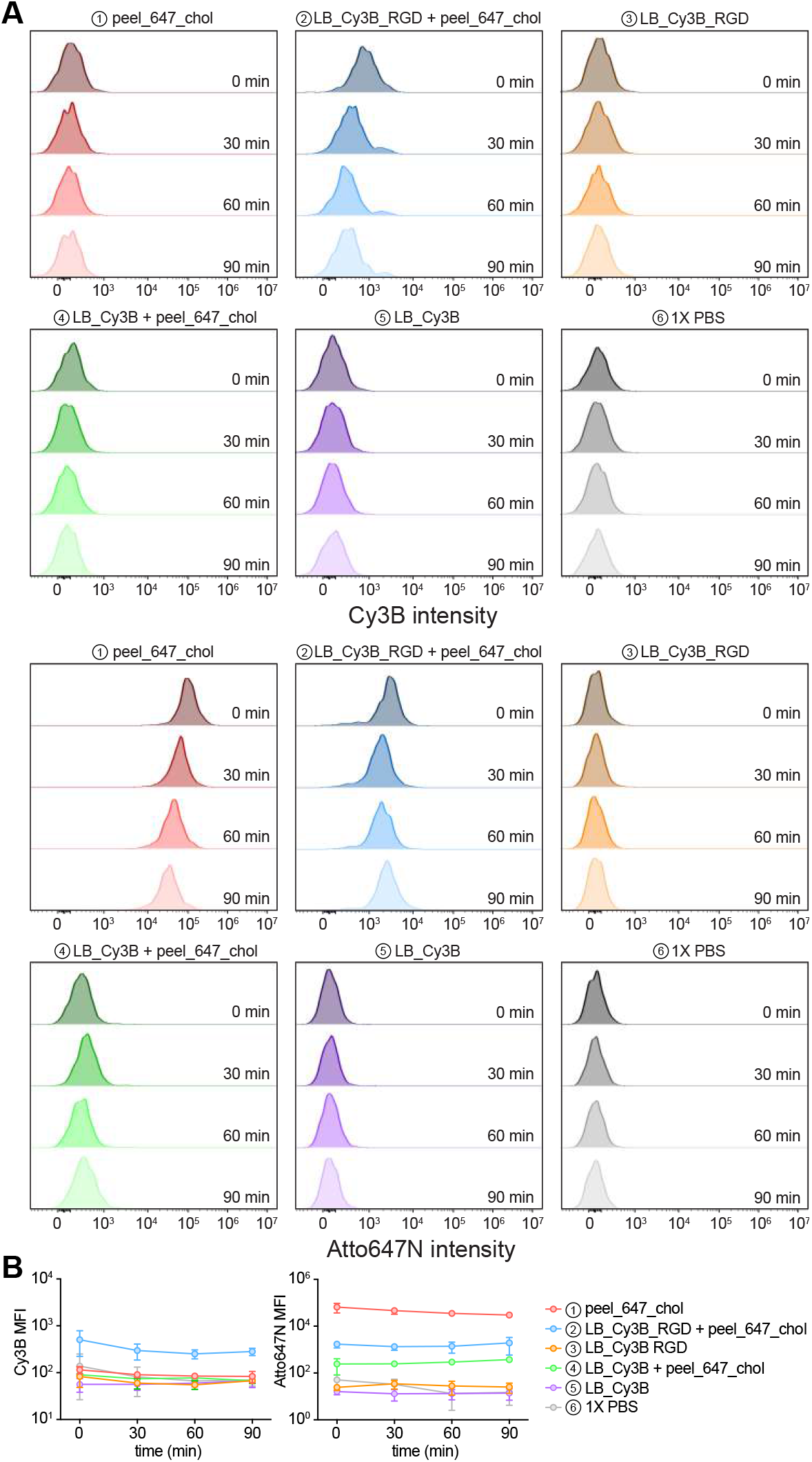
Cholesterol-DNA incorporation in mouse platelets. Mouse platelets were incubated with 50 nM of different DNA strands and duplexes with or without cholesterol and RGD in Tyrode’s buffer at room temperature for 30 min as described in **Extended Figure 4 (see Extended Figure 4A for scheme describing different structures)**. After incubation, platelets were spun down, washed twice, and resuspended in Tyrode’s buffer. Samples were divided into 4 portions and the Cy3B and Atto647N fluorescence in cells was measured at t = 0 min, 30 min, 60 min, and 90 min. (A) Representative flow cytometry histogram showing the Cy3B and Atto647N fluorescence in platelets after incubation with DNA strands and duplexes at different time points. (B) Plot showing the Cy3B and Atto647N MFI of platelets incubated with different DNA strands and duplexes from n = 3 experiments.

**Figure S7.**
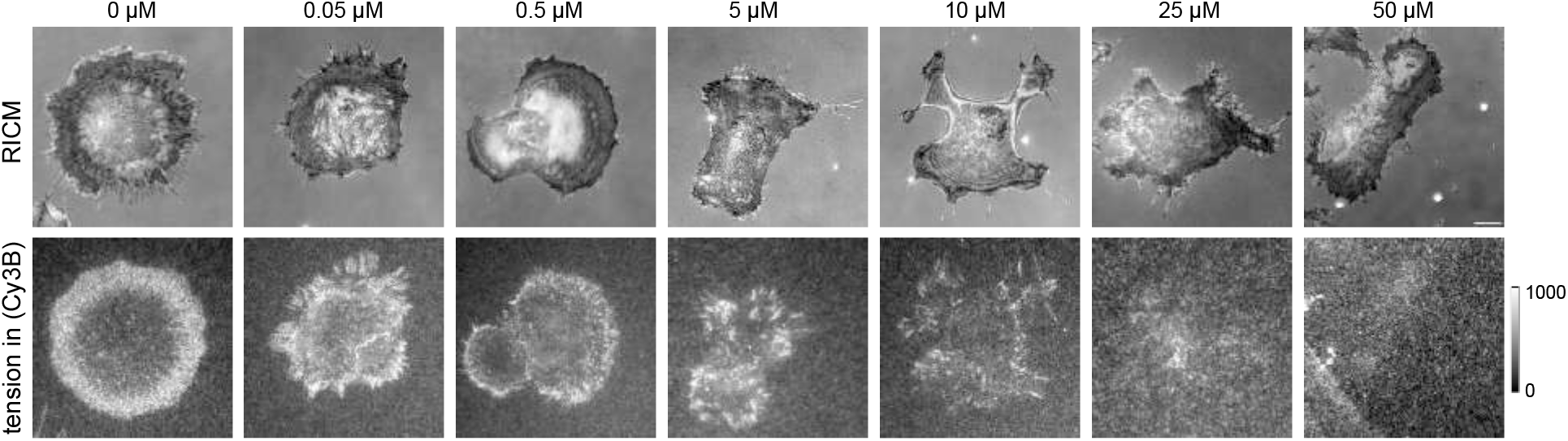
Representative microscopy images of cells pretreated with Y27632. RICM images show the spreading of the cells pretreated with Y27632 at t = 60 min. The tension signal in Cy3B channel show the irreversible tension over 60 min after plating the pretreated cells on peeling probe substrates. Scale bar = 10 µm.

**Figure S8.**
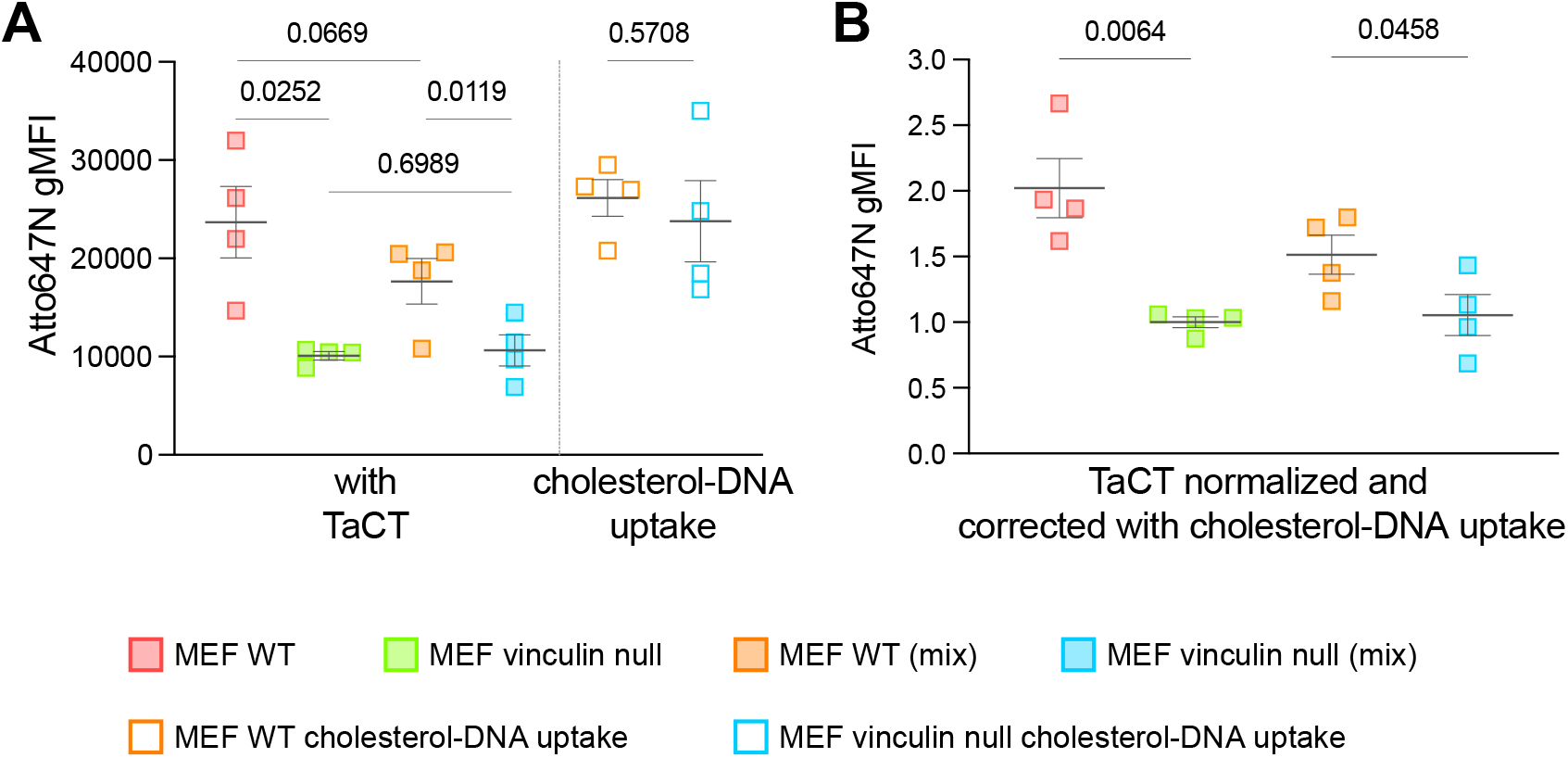
TaCT assay results in MEF WT and MEF vin-cells. (A) Plot shows the Atto647N gMFI of MEF WT and MEF vin-cells after TaCT when cells were seeded on separate TaCT substrates or mixed and seeded on the same TaCT substrate as described in the maintext and **Figure 2**. The cholesterol-DNA incorporation was measured as a reference for potential differential uptake in WT and vin-cells. WT and vin-cells were incubated with 10 nM Atto647N labeled cholesterol peeling strand in solution in parallel to TaCT assays and the incorporation was measured with the flow cytometer. We chose 10 nM as the cholesterol uptake control, based on microscopy analysis indicating that on average ∼46790±2983 peeling strands (7.772×10^™20^ mol) were released per fibroblast. Assuming the height of a spreading cell (mean spreading area = ∼1280 µm^2^) is ∼5 µm, the estimated volume around the cell would be ∼6.4×10^™12^ L, and the local concentration of the released cholesterol-DNA would be ∼12 nM. (B) Plot shows the Atto647N gMFI values normalized to the cholesterol-DNA uptake. Data acquired from 4 replicates, and statistical analysis was performed using paired Student t-test.

**Figure S9.**
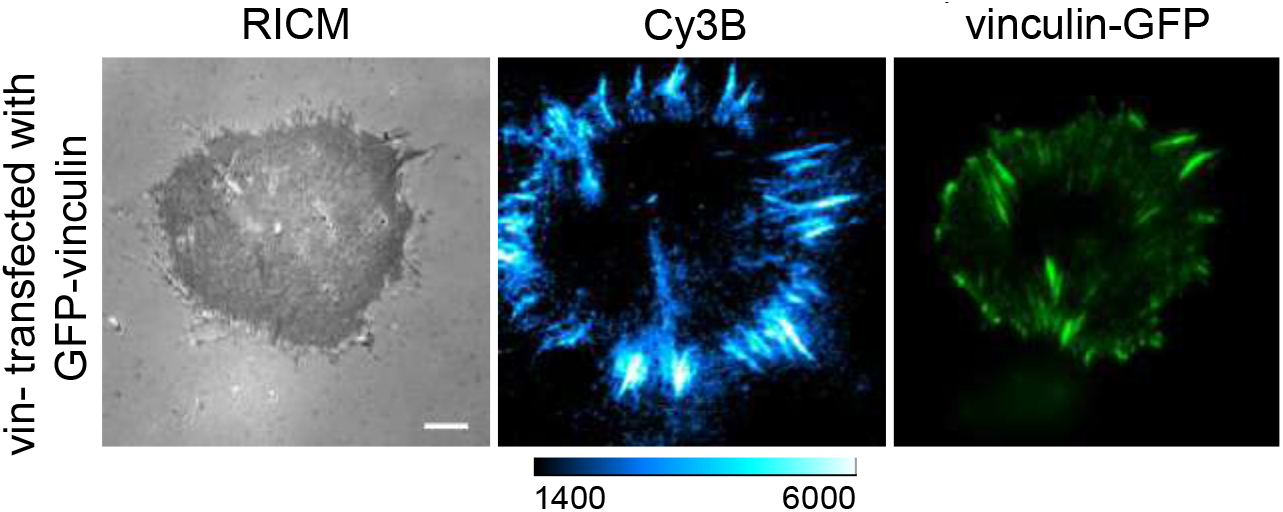
Representative microscopy images of a MEF vin-cell transfected with 3 µg of GFP-vinculin. Cells were transfected with GFP-vinculin plasmid for 24 h, and then seeded onto TaCT substrates for integrin tension measurements. Images of cell spreading, integrin tension, and vinculin expression were acquired at t=60 min. Scale bar = 10 µm.

**Figure S10.**
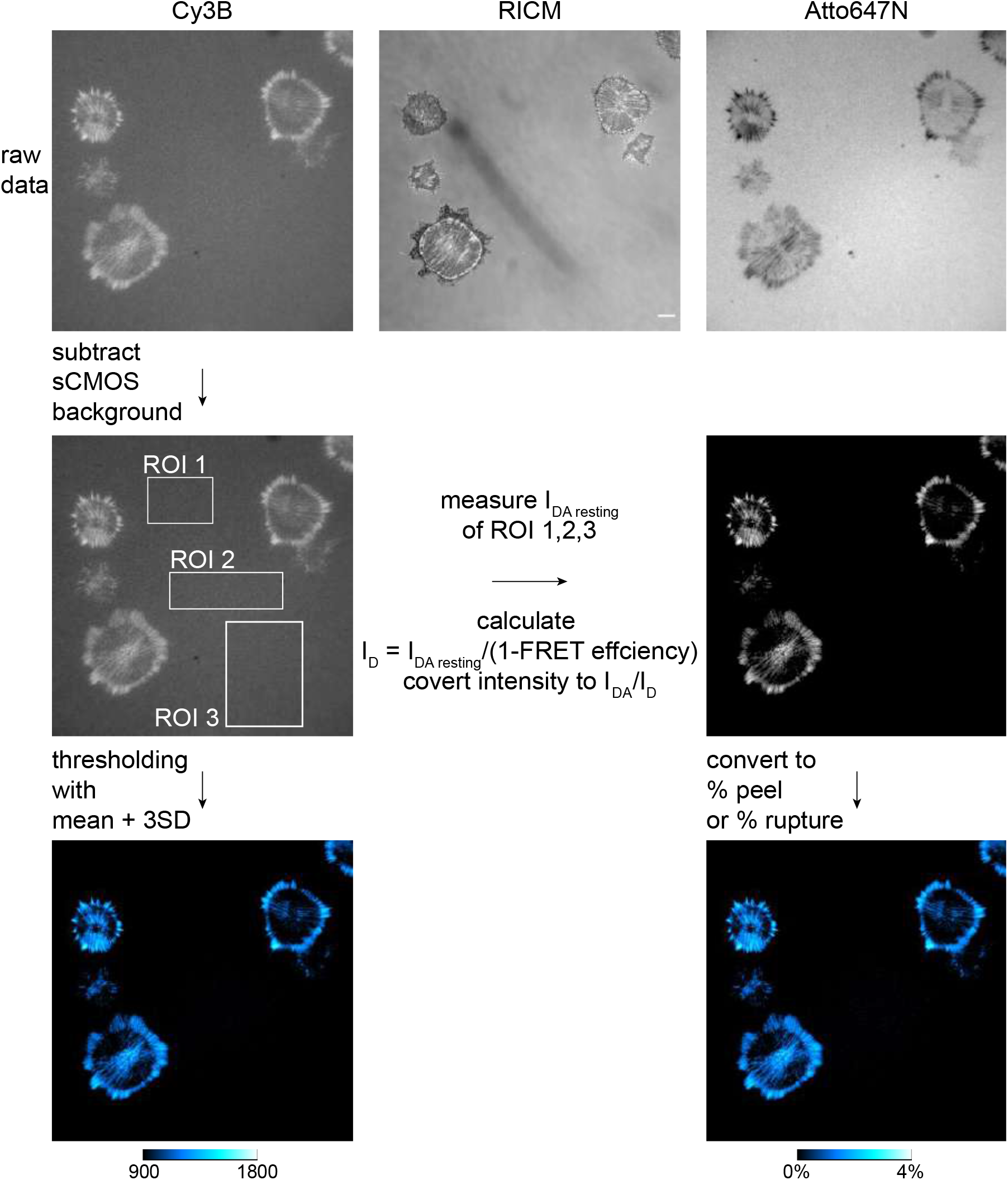
Microscopy data analysis workflow. Raw fluorescence imaging data was collected and the sCMOS background was subtracted. Three local ROIs were drawn, and the duplex probe background I_DA resting_ was measured and averaged. Then, the fluorescence of the fully peeled background I_D_ was calculated using FRET efficiency calculated from **Figure S4A**. The image was then divided by I_D_ to obtain an I_DA_/I_D_ tension image. The I_DA_/I_D_ image was then converted to %peel by applying a predefined conversion function in **Figure S4B, C**.

**Figure S11.**
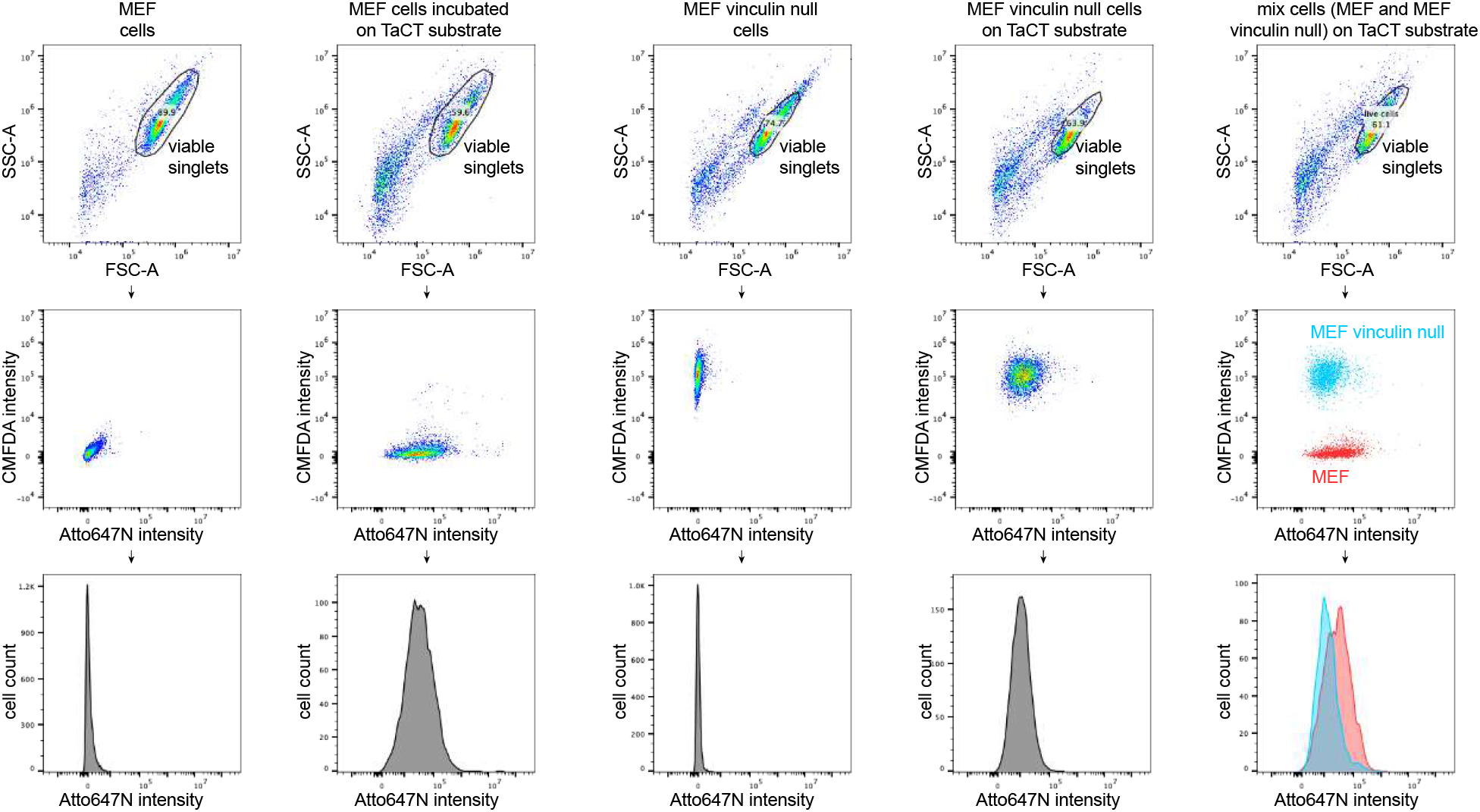
Representative flow cytometry data analysis. For the flow cytometry analysis of TaCT, viable cells and singlets cells were identified using the forward scatter and side scatter, and the histogram was used to show the fluorescence intensity of viable singlets.

## 4. Videos

**Video S1. Simulation of 24 bp DNA peeling with oxDNA**.

**Video S2. Fibroblast cell producing integrin tension > 41 pN**.

## Notes

### Competing Interest Statement

The authors have declared no competing interest.

